# Adolescent maturation of cortical excitation-inhibition balance based on individualized biophysical network modeling

**DOI:** 10.1101/2024.06.18.599509

**Authors:** Amin Saberi, Kevin J. Wischnewski, Kyesam Jung, Leon D. Lotter, H. Lina Schaare, Tobias Banaschewski, Gareth J. Barker, Arun L.W. Bokde, Sylvane Desrivières, Herta Flor, Antoine Grigis, Hugh Garavan, Penny Gowland, Andreas Heinz, Rüdiger Brühl, Jean-Luc Martinot, Marie-Laure Paillère Martinot, Eric Artiges, Frauke Nees, Dimitri Papadopoulos Orfanos, Herve Lemaitre, Luise Poustka, Sarah Hohmann, Nathalie Holz, Christian Baeuchl, Michael N. Smolka, Nilakshi Vaidya, Henrik Walter, Robert Whelan, Gunter Schumann, IMAGEN Consortium, Tomáš Paus, Juergen Dukart, Boris C. Bernhardt, Oleksandr V. Popovych, Simon B. Eickhoff, Sofie L. Valk

## Abstract

The balance of excitation and inhibition is a key functional property of cortical microcircuits which changes through the lifespan. Adolescence is considered a crucial period for the maturation of excitation-inhibition balance. This has been primarily observed in animal studies, yet human *in vivo* evidence on adolescent maturation of the excitation-inhibition balance at the individual level is limited. Here, we developed an individualized *in vivo* marker of regional excitation-inhibition balance in human adolescents, estimated using large-scale simulations of biophysical network models fitted to resting-state functional magnetic resonance imaging data from two independent cross-sectional (N = 752) and longitudinal (N = 149) cohorts. We found a widespread relative increase of inhibition in association cortices paralleled by a relative age-related increase of excitation, or lack of change, in sensorimotor areas across both datasets. This developmental pattern co-aligned with multiscale markers of sensorimotor-association differentiation. The spatial pattern of excitation-inhibition development in adolescence was robust to inter-individual variability of structural connectomes and modeling configurations. Notably, we found that alternative simulation-based markers of excitation-inhibition balance show a variable sensitivity to maturational change. Taken together, our study highlights an increase of inhibition during adolescence in association areas using cross sectional and longitudinal data, and provides a robust computational framework to estimate microcircuit maturation *in vivo* at the individual level.

## Introduction

The vast repertoire of cortical functions emerges from a careful tuning of the interactions between excitatory (E) and inhibitory (I) neurons in microcircuits embedded in the structural scaffolding of the brain^1^. Excitation and inhibition, mainly transmitted via glutamate and γ-aminobutyric acid (GABA), respectively, are inseparable and balanced, that is, the inhibition generated in the cortical microcircuits is proportional to the local and incoming excitation^2^. This phenomenon has been observed during both responses to external stimuli^3–5^ and spontaneous cortical activity^4,6^. The E-I balance is proposed to be essential for central aspects of cortical functioning, including the dynamic stability of activity^7^, efficient coding of the information^8^, sharp tuning of sensory stimuli^2^, and generation of synchronous cortical oscillations in gamma and beta ranges^2,6,9,10^. Conversely, disturbed E-I balance can lead to cortical circuit dysfunctioning and is hypothesized as a key pathophysiological mechanism in various neuropsychiatric conditions such as schizophrenia, autism spectrum disorder, and epilepsy^11–15^.

Adolescence is a critical developmental period with substantial changes in the brain including maturation of the E-I balance^16–18^. During this period, several important changes occur in the architecture and function of excitatory and inhibitory neurons and synapses, which together are suggested to lead to a re-calibration of the E-I balance^16^. For instance, *post-mortem* histology of adolescent brains has shown a pruning of excitatory synapses within the prefrontal cortex, in rats^19,20^, non-human primates^21,22^ and humans^23–25^. In addition, *post-mortem* transcriptomic studies of the prefrontal cortex in animals and humans have indicated marked changes in the expression of genes involved in inhibitory neurons and GABAergic signaling, including parvalbumin^9,26,27^ and GABA^A^ receptor subunits^28–31^. These transcriptomic changes are accompanied by the maturation of inhibitory function with stronger and shorter inhibitory postsynaptic currents, as observed in the prefrontal cortex of non-human primates^9,30^, overall indicating a relative increase of inhibitory synaptic transmission in this area^16,32^.

Currently available evidence on the *in vivo* maturation of the E-I balance in humans is limited, as the invasive methods used in animal studies are not feasible in humans. However, *in vivo* proxies of E-I balance have been proposed, relying on its putative macroscale functional consequences captured in functional imaging^33,34^ and electrophysiology^35–37^, or through biochemical quantification of glutamatergic or GABAergic neurotransmitters using magnetic resonance spectroscopy^38–40^. Such approaches are informative, but lack a certain level of mechanistic insight and detail that is observed with, for example, direct measurement of the excitatory and inhibitory input currents as done in animal research. Furthermore, studies on the development of E-I balance are often focused on selected areas, primarily within the prefrontal cortex, and the knowledge on the regional patterns of E-I balance maturation across the whole cerebral cortex is limited. Biophysical network modeling (BNM) of the brain is a promising computational technique that can bridge different scales of investigation at a whole-cortical level. It provides a tool to non-invasively derive mechanistic inferences about a hidden brain feature at the microscale, such as the E-I balance, based on the observed (empirical) *in vivo* data at the macroscale, and has provided valuable insights into brain (dys)function^41–47^. In this approach, dynamic spontaneous activity of brain areas are simulated using biologically realistic models which are informed by, for instance, the blood-oxygen-level-dependent (BOLD) signal measured during resting-state functional magnetic resonance imaging (rs-fMRI)^48–51^.

In this study, we aimed to investigate the adolescent maturation of the regional E-I balance in humans at the individual level, by applying the BNM approach on two independent cross-sectional and longitudinal neuroimaging datasets from the Philadelphia Neurodevelopmental Cohort (PNC) and the IMAGEN study^52,53^. We performed large-scale simulations of individualized BNMs^44,54,55^, in which models were informed by structural connectivity (SC) and functional imaging data of each subject. The subject-level precision of these models allowed for mapping the estimated E-I balance specifically in each individual using simulations that best represented their empirical data, and furthermore enabled studying within-subject maturation longitudinally. By doing so, we extended a previous study that used the BNM approach to study the development of the E-I balance in the PNC dataset at the level of age groups^56^. We demonstrated replicable effects across the two datasets indicating cross-sectional and longitudinal age-related increases of relative inhibition in the association areas and a lack of significant changes or even relative increases of excitation in the sensorimotor areas. We found this neurodevelopmental pattern of the shifts in E-I balance was aligned with the proposed sensorimotor-association axis of the cortical neurodevelopment^18^. Subsequently, given that the simulation results may be affected by various modeling and analytical choices^57–59^, or might be confounded by the variability of underlying structural connectome, as well as the noise within the simulations and parameter optimization, we extensively assessed and demonstrated the robustness of our simulation-based findings against these nuisances. Lastly, we contrasted our marker of the E-I balance to alternative previously-used BNM-based markers and highlight methodological and conceptual considerations regarding the usage and interpretation of these markers.

## Results

### Overview

We included 752 adolescents from the cross-sectional PNC dataset^52^ (409 female, mean age: 15.3±2.4 [10-19] years) and 149 participants from the longitudinal IMAGEN study^53^ (72 female, mean age: 14.4±0.4 years at baseline, 18.9±0.5 years at follow-up). Subject/session diffusion-weighted imaging (DWI) and rs-fMRI data was used to generate individual matrices of: i) structural connectome based on the density of white matter streamlines, ii) functional connectivity (FC) matrix as the correlation of the BOLD signals, and iii) functional connectivity dynamics (FCD) matrix as a measure of how the FC dynamically evolves through sliding windows of time during the scan, across 100 cortical areas^60^. Hereafter, we refer to FC and FCD matrices derived from the imaging data as *empirical FC and FCD* to distinguish them from *simulated FC and FCD*.

Next, we performed individualized BNM simulations and parameter optimizations for each subject/session to estimate their *in silico* regional measures of the E-I balance based on their *in vivo* imaging data (Figure 1). We applied the reduced Wong-Wang model^61^, which models each node as coupled excitatory and inhibitory neuronal pools, where the excitatory neuronal pools of different nodes are interconnected through the individual-specific SC. The model was controlled by global and regional free parameters which were fit to the empirical resting-state functional data of the target subject/session using the covariance matrix adaptation-evolution strategy (CMA-ES) optimization algorithm^62–64^. This involved running a maximum of 33600 simulations per subject/session using an efficient implementation of BNM simulations on graphical processing units (GPUs; https://cubnm.readthedocs.io). The model parameters included a global parameter *G*which scales the strength of inter-regional coupling, in addition to regional parameters 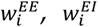 and 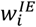 which characterize the connectivity weights between E and I neuronal pools within each node. Motivated by recent developments of this model^65,66^, we let 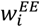 and 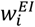 to vary across nodes, i.e., they were computed, independent of each other, through weighted combinations of six fixed biological maps that represent morphological, functional, genetic and neurochemical heterogeneity of the human cerebral cortex. These maps were obtained from independent healthy adult samples, and included average T1-weighted/T2-weighted ratio (T1w/T2w), average cortical thickness, principal gradient of FC (FC G1), principal axis of gene expression (Gene PC1), and average NMDA and GABA^A/BZ^ receptor PET maps^67–75^ (Figure S1). Furthermore, in each simulation 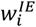 was determined based on an analytical-numerical feedback inhibition control (FIC) algorithm, which aimed to maintain the firing rate of excitatory neurons within a biologically plausible range of 3 Hz^61,66^. Following, from the optimal simulations of each subject/session we extracted the *in silico* input current to the excitatory neuron of each node 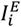, averaged across simulation time, which resulted in individual-specific 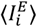 maps. The 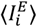 values reflect *in silico* estimates of regional E-I balance, given 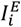 results from the combination of excitatory input currents to each node (from itself, and from the excitatory neurons of the other nodes through the SC) balanced by local inhibitory currents. Therefore, increase of 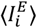 can be interpreted as a relative increase of excitation or decrease of inhibition within a model region.

**Figure 1.**
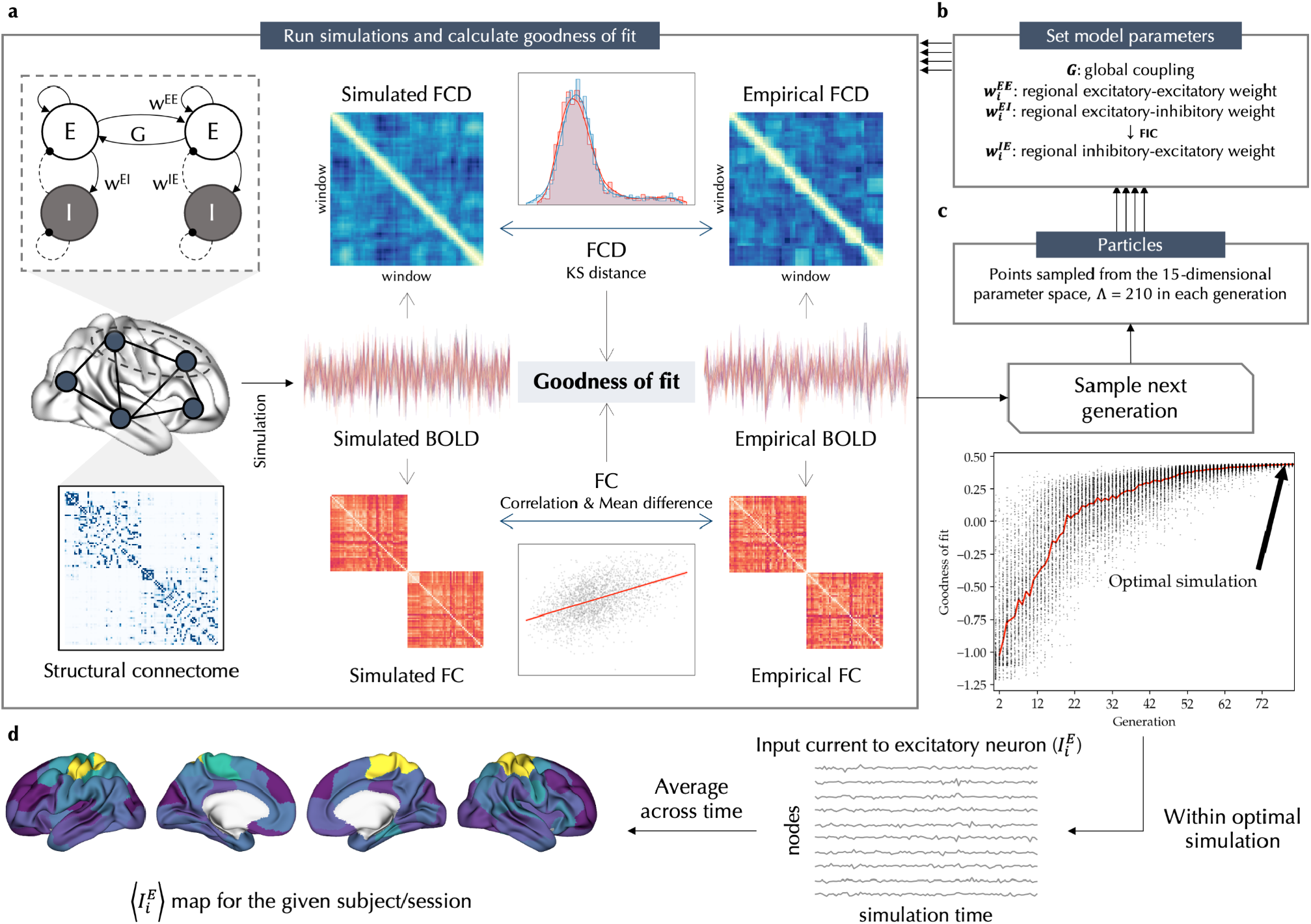
Overview. Individualized biophysical network model (BNM) simulation-optimization (**a-c**) was performed to derive the subject-/session-specific regional measures of the E-I balance, defined as time-averaged *in silico* input current to the excitatory neurons, 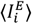 (**d**). The model consists of coupled excitatory (E) and inhibitory (I) neuronal pools in each node, where the E neuronal pools of brain nodes are interconnected through the structural connectome (SC) of the given subject/session (**a**; *left*). The model is controlled by a global parameter *G* which adjusts inter-regional coupling, in addition to regional parameters 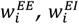 and 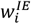 which characterize the connection weights between E and I neuronal pools within each node. In each simulation, *G*, 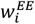 and 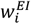 are set by the optimizer while 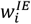 is determined based on feedback inhibition control (FIC) algorithm (**b**). The covariance matrix adaptation-evolution strategy (CMA-ES) algorithm was used to optimize model parameters with respect to empirical functional data of the given subject (**c**). The optimization goal was to maximize the goodness-of-fit by tuning 15 free parameters, including *G*, as well as bias and coefficient terms used to determine 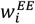 and 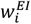 based on six fixed biological cortical maps (Figure S1). In each generation of the optimizer, 210 simulations (“particles”) with different parameters were performed. The goodness-of-fit of each simulation to the empirical functional data (**a**, *right*) is assessed as the correlation of functional connectivity (FC) matrices subtracted by their absolute mean difference and the Kolmogorov-Smirnov distance of functional connectivity dynamics (FCD) matrices derived from the simulated or empirical blood-oxygen-level-dependent (BOLD) signal. After completion of two CMA-ES runs, each with a maximum of 80 generations, the optimal simulation with the best goodness-of-fit to the empirical functional data of the target subject/session is selected (**c**). Lastly, the *in silico* input current to the excitatory neuron of each node 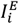 is averaged across simulation time, resulting in an 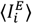 map for each subject/session, which was used as a regional marker of the excitation-inhibition balance in this study (**d**).

### Cross-sectional age-related variation of the E-I balance

Following, we studied cross-sectional age-related variation of the excitation-inhibition balance in adolescence, estimated based on 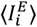, in the PNC dataset. The individualized optimal simulations of the PNC dataset showed a goodness-of-fit of 0.259±0.101 to the empirical data (Figure S2). Based on these simulations, we found widespread significant age-related decreases of 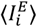 in association areas within the frontal, parietal, and temporal lobes, in contrast to its age-related increases in visual and sensorimotor areas as well as the left posterior insula, after controlling for goodness-of-fit, sex, and in-scanner rs-fMRI motion, and adjusting for multiple comparisons at a false discovery rate (FDR) of 5% (Figure 2a). The effect sizes across these regions, calculated as Pearson correlation coefficient between age and 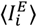 controlled for the confounds, ranged between -0.255 to 0.156. We then assessed within-sample stability of the age effects across 100 subsamples of the data, each including half of the total sample with 376 subjects. The unthresholded age effects on 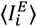 across all pairs of subsamples were correlated with a mean correlation coefficient of 0.863±0.047, indicating high within-sample stability of the observed age effects (Figure 2c). Of note, assessing maturational differences of 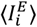 between males and females, we found no significant age-by-sex interactions after FDR correction.

**Figure 2.**
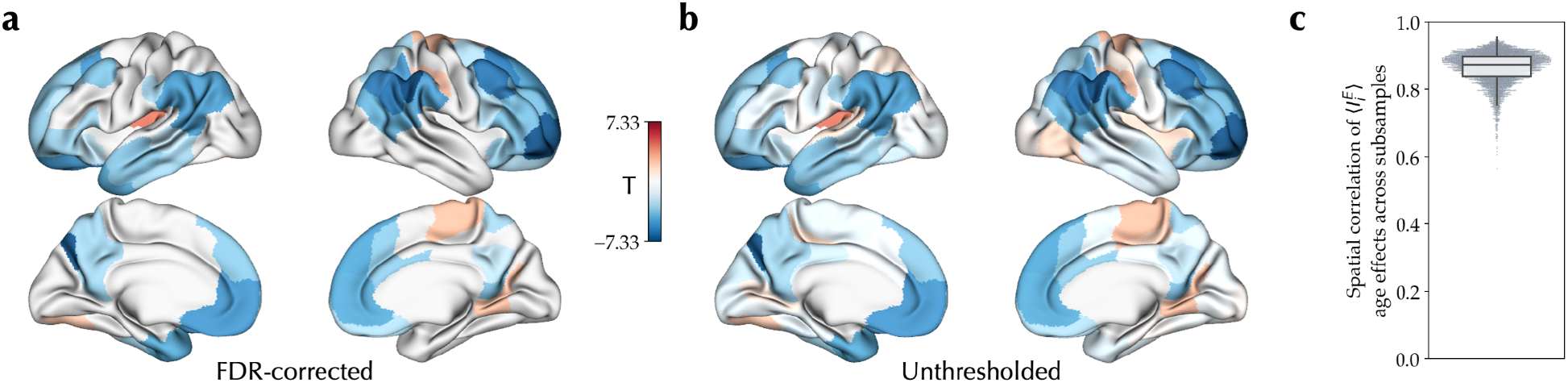
Cross-sectional effect of age on excitation-inhibition balance during adolescence. **(a)** Linear effect of age on 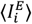, showing its significant age-related decrease (blue) and increase (red) during adolescence in the PNC dataset, after removing outliers and controlling for goodness-of-fit, sex and in-scanner rs-fMRI motion, corrected for multiple comparisons using false discovery rate (FDR). **(b)** Unthresholded map of the effect of age on 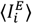. **(c)** The distribution of correlation coefficients between 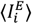 age effect maps of all pairs of subsamples across 100 half-split subsamples of the dataset.

### Longitudinal changes of the E-I balance

To extend and assess the replicability of our findings in the cross-sectional PNC study, we next investigated the longitudinal E-I maturation in the independent IMAGEN dataset, including 149 participants assessed at the ages of 14 and 19 years. The individualized optimal simulations in the IMAGEN had a mean goodness-of-fit of 0.266±0.102 at the baseline and 0.231±0.113 at the follow-up session (Figure S3). Within these simulations, we found a significant longitudinal age-related decrease of 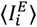 in widespread association areas within the frontal, parietal and temporal lobe, and a significant increase in visual areas, controlled for goodness-of-fit, sex, in-scanner rs-fMRI motion, and site, and adjusted for multiple comparisons at FDR of 5% (Figure 3a). The effect sizes across these regions, calculated as the standardized mean difference of baseline to follow-up session in 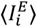 controlled for the confounds, ranged between -0.299 to 0.229. We next assessed the within-sample stability of age effects using 100 subsamples of the IMAGEN data, each including half of the total sample with 74 subjects. This resulted in a mean correlation of r = 0.746±0.108 between the 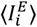 age effect maps across all pairs of subsamples (Figure 3c). Furthermore, we found no significant age-by-sex interactions after FDR correction, yielding no evidence for sex differences in the maturation of 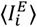. Of note, given the quality of the tractograms in the baseline session of the IMAGEN dataset was lower, in these simulations we used the SC of the follow-up session in the models of both sessions. However, in a subset of subjects with adequate quality of tractograms in both baseline and follow-up sessions (N = 110; 52 female), using models with session-specific SCs resulted in largely similar effects of age on 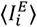 (r = 0.779, p_spin_ < 0.001; Figure S4).

**Figure 3.**
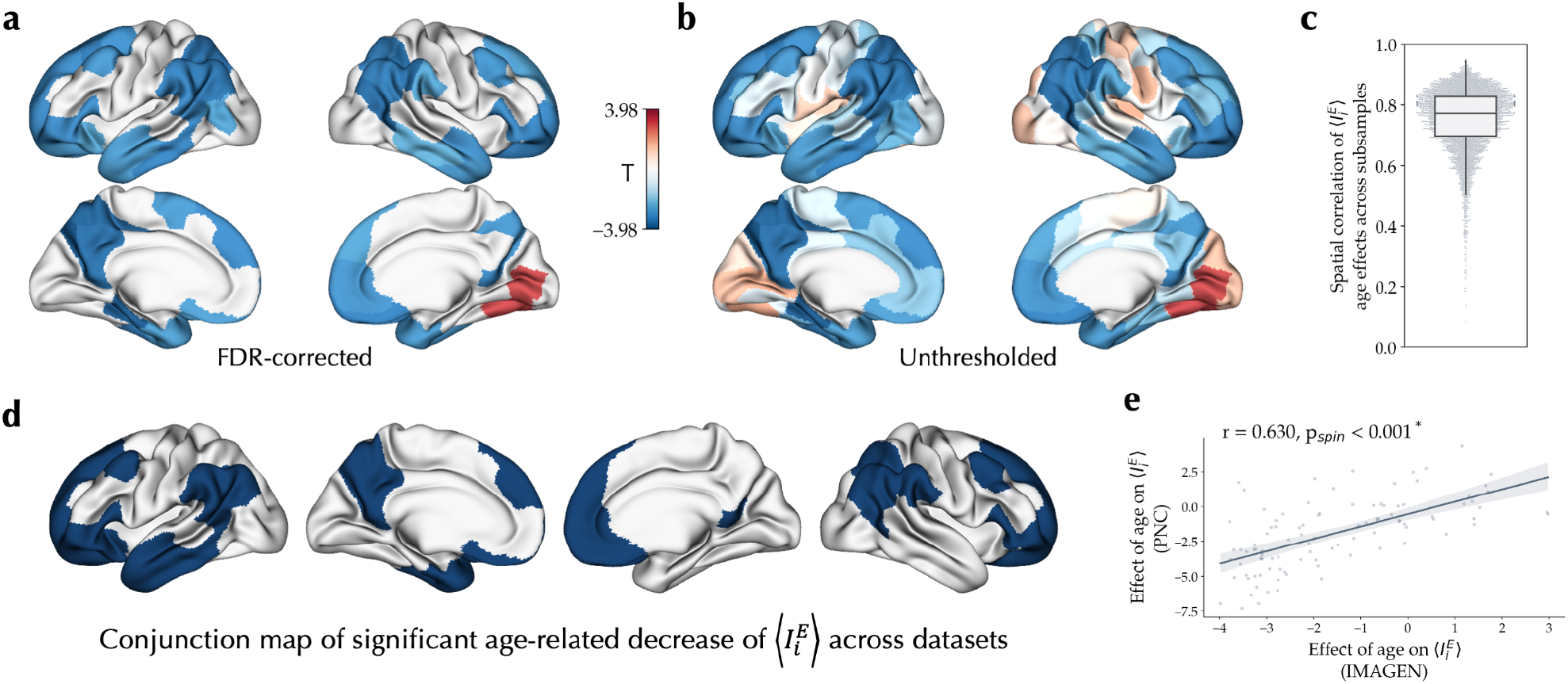
Longitudinal effect of age on the excitation-inhibition balance during adolescence. **(a)** Linear effect of age on 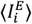, showing its significant longitudinal decrease (blue) and increase (red) through adolescence, using a mixed effects model with random intercepts for each subject, after removing outliers and controlling for goodness-of-fit, sex, in-scanner rs-fMRI motion and site, and corrected for multiple comparisons using false discovery rate (FDR). **(b)** Unthresholded effect of age on 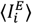. **(c)** The distribution of correlation coefficients between 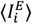 age effect maps of all pairs of subsamples across 100 half-split subsamples of the dataset. **(d)** Conjunction of regions showing significant decreases of 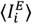 associated with age in the PNC and IMAGEN datasets. **(e)** Spatial correlation of longitudinal effects of age on 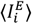 in IMAGEN with cross-sectional effect of age on 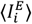 in PNC.

We next assessed the similarity of cross-sectional and longitudinal age-related variation of 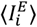 observed in the two datasets. Conjunction of regions with significant age effects on 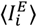 in the PNC and IMAGEN datasets revealed 33 regions in the association cortices showing significant decrease of 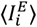, whereas no region showed significant replicable age-related increase of 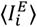 (Figure 3d). Mean 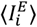 across these association regions, after regressing out the effects of confounds, showed a correlation coefficient of r = -0.232 with age in the PNC (T = -6.64, p < 0.001), and a standardized mean difference of -0.312 between the sessions in IMAGEN (T = -4.17, p < 0.001; Figure S5). Furthermore, the unthresholded map of 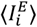 longitudinal age effects in IMAGEN (Figure 3b) was significantly correlated (r = 0.630, p_spin_ < 0.001; Figure 3e) with the cross-sectional age effect map observed in PNC (Figure 2b).

Therefore, overall, across the two datasets we observed replicable cross-sectional and longitudinal effects indicating a developmental relative increase of inhibition in the association areas in contrast to a lack of significant changes or a relative increase of excitation in sensorimotor areas.

### Neurodevelopmental pattern of EI balance co-aligns with the sensorimotor-association axis of cortical organization

Having observed differential effects of age on the E-I balance across cortical areas, we next sought to investigate the embedding of this spatial neurodevelopmental pattern across different domains of cortical organization as well as meta-analytic maps of cortical function and developmental transcriptomics.

We first studied the spatial co-alignment of the observed age effects on 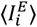 with a previously proposed sensorimotor-association axis of cortical neurodevelopment and the multimodal cortical features it was composed of^18^ (Figure S6, Table S1). The 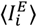 age effect maps of both datasets were significantly (p_spin_ < 0.001) correlated with the sensorimotor-association axis map (PNC: r = -0.617; IMAGEN: r = -0.607) as well as several of its components, notably including FC G1 (PNC: r = -0.691; IMAGEN: r = -0.641) and T1w/T2w (PNC: r = 0.437; IMAGEN: r = 0.548; Figure 4a, S7a). Next, comparing the 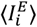 age effects across seven canonical resting-state networks^76^, we observed more negative age effects in the default mode and frontoparietal networks compared to the somatomotor and visual networks (Figure 4b, S7b). These findings indicated co-alignment of the E-I balance maturational pattern with the sensorimotor-association axis of the cortex with higher age-related relative increase of inhibition towards the association areas.

**Figure 4.**
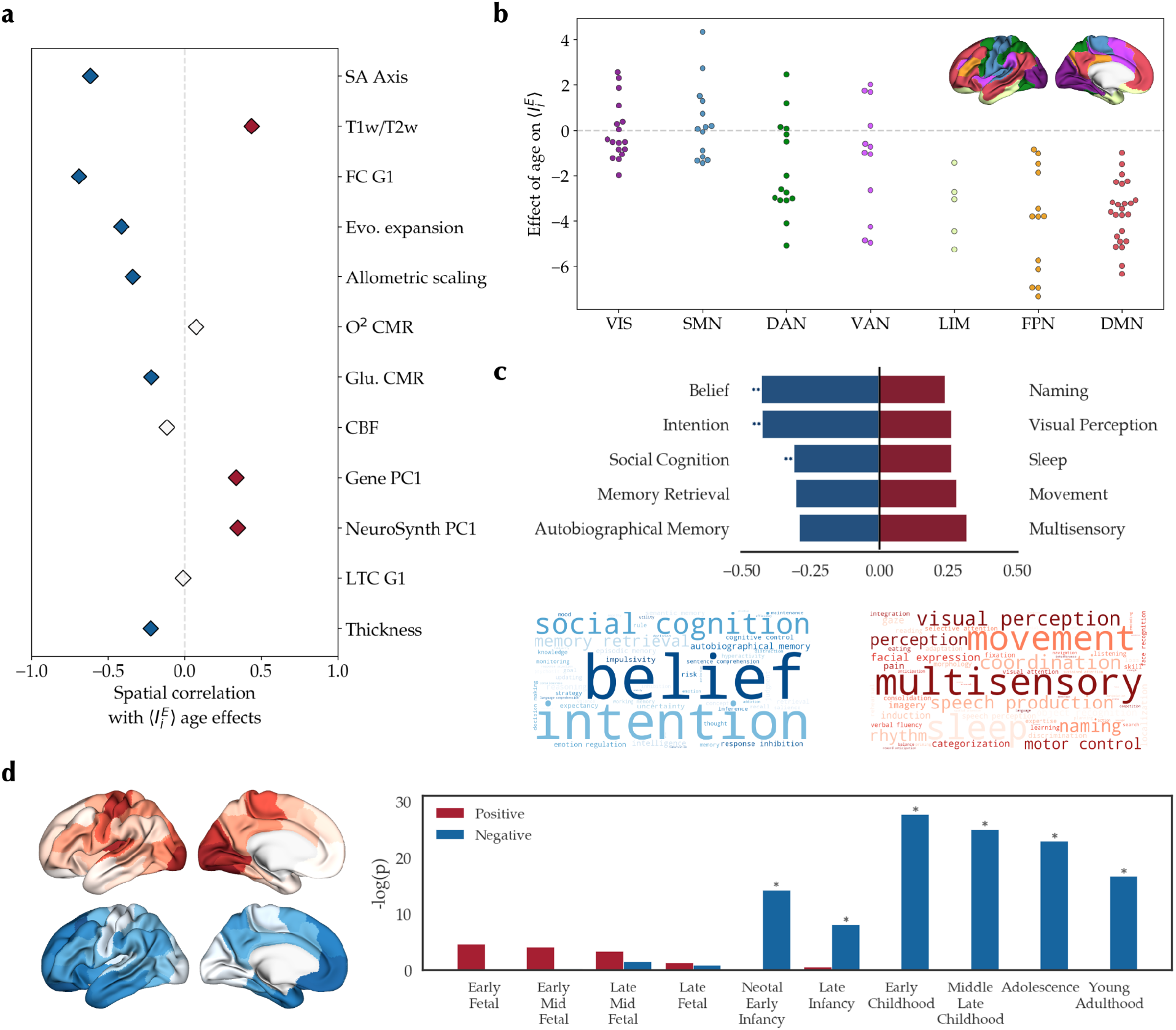
Embedding of the excitation-inhibition developmental pattern in the PNC dataset along the sensorimotor-association axis. (**a**) Spatial correlation of 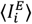 age effect in the PNC dataset with maps of sensorimotor-association cortical axis based on Sydnor et al.^18^ (Figure S6). Colored diamonds show statistically significant (p_spin_ < 0.05) positive (red) and negative (blue) spatial correlations. **(b)** Distribution of 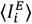 age effects across the canonical resting-state networks (F = 13.85, p_spin_ < 0.001). Post-hoc tests (Bonferroni-corrected) showed significantly more positive age effects in visual (VIS) and somatomotor (SMN) compared to limbic (LIM), frontoparietal (FPN) and default mode networks (DMN), in addition to more positive age effects in dorsal attention network (DAN) compared to DMN. **(c)** *top*: Meta-analytical maps with significant spin correlation to the 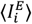 age effect map. Double asterisks denote associations that were significant after false discovery rate (FDR) adjustment. *bottom*: Word clouds of meta-analytical maps negatively (blue) or positively (red) correlated with the 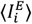 age effect map. Size of the words is weighted by their correlation coefficient. **(d)** *left*: Mean expression of the top 500 genes associated with the 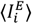 age effect map split into sets of negatively-(N = 187, blue) and positively-associated (N = 313, red) genes. *right:* Specific expression analysis of the two sets of genes across developmental stages in the cortex. Y-axis shows the negative log of uncorrected p-values. Asterisks denote significantly enriched developmental stages after FDR adjustment. ===== SA: sensorimotor-association, T1w/T2w: T1-weighted to T2-weighted ratio, FC G1: principal gradient of functional connectivity, Evo.: evolutionary, CMR: cerebral metabolic rate, Glu.: glucose, CBF: cerebral blood flow, Gene PC1: principal axis of Allen Human Brain Atlas gene expression data, NeuroSynth PC1: Principal component of NeuroSynth meta-analytical maps, LTC G1: principal gradient of laminar thickness covariance.

Following, we assessed the functional relevance of the observed developmental patterns by evaluating their spatial correlation with meta-analytical maps of terms related to cognitive and behavioral functions^70^ obtained from the NeruoSynth database^77,78^ (123 terms, Table S2). We found significant (p_spin_ < 0.001) negative correlations of the age effect maps with the maps of terms such as “intention” (PNC: r = -0.425; IMAGEN: r = -0.376) and their positive correlations with the maps of terms such as “visual perception” (PNC: r = 0.261; IMAGEN: r = 0.256), overall suggesting that areas which show a relative increase of inhibition during adolescence are associated with higher-order functions (Figure 4c, S7c).

Lastly, we performed developmental transcriptomics enrichment analysis of 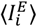 age effect maps. Using partial least squares regression with the gene expression maps obtained from the Allen Human Brain Atlas^71,73^, we identified the top 500 genes expressed higher towards the negative and positive ends of 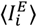 age effect maps. Next, we investigated the developmental enrichment of the two sets of genes using specific expression analysis of BrainSpan dataset^79^ and found that the genes expressed towards the negative ends of 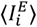 age effect maps to be enriched in later stages of development, primarily during and after childhood, in contrast to the genes expressed towards the positive ends of 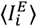 age effect maps that were enriched in earlier fetal stages of development, though not significantly (Figure 4d, S7d).

### Robustness analyses

Thus far, we observed consistent age effects in the adolescent maturation of E-I balance in two independent cross-sectional and longitudinal datasets across a sensorimotor-association axis by using simulations of individualized BNMs. However, these simulation-based findings may be sensitive to various modeling and analytical choices^57–59^ as well as confounding effects of the underlying structural connectome or noise. Therefore, we next assessed the robustness of 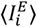 and its age-related changes to such nuisances, including the effects of inter-individual variability of SC, modeling configurations, and the randomness within the optimizer and simulations. For brevity and to reduce the usage of computational resources, we limited these analyses to the PNC dataset.

#### Inter-individual variability of structural connectome

Using subject-specific SCs in the main analyses enabled modeling of brain function within an individualized structural scaffold which better represents each subject. However, this potentially introduces inter-individual variability of SCs as a source of variability in 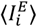, particularly given 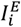 is directly related to the SC in the model Eq. 1. As a result, the associations of age with model-derived features may be confounded by age-related variation of trivial features of SC, such as the node-wise strength. However, the age effect on 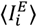 was robust to controlling for the node-wise SC strength as an additional confound (r = 0.959, p_spin_ < 0.001; Figure 5c, Figure S8a). Furthermore, when SC variability was eliminated by using an identical template SC in the BNM simulations of all the subjects, we observed an age effect on 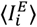 largely consistent with the effects observed in main analyses (r = 0.618, p_spin_ < 0.001; Figure 5c, S8b).

**Figure 5.**
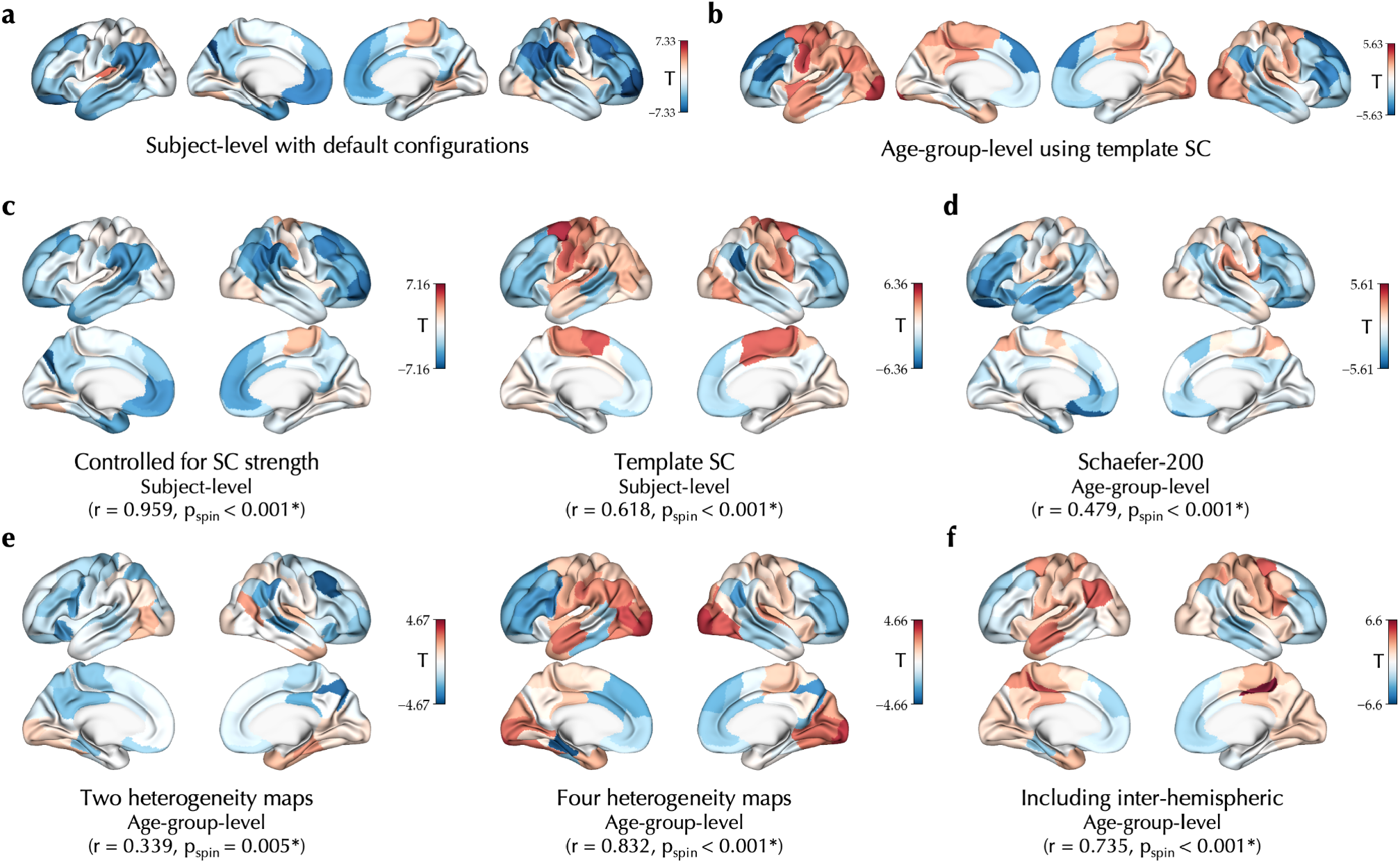
Robustness analyses. The unthresholded effect of age on 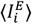 observed in the PNC dataset using the default configurations at subject level (**a**) or age-group level using template SC (**b**) compared to age effects observed using alternative configurations, including: **(c**, *left*) linear regression additionally controlled for the node-wise structural connectome (SC) strength, that is, row-wise sum of the SC matrix, **(c**, *right*) using a fixed template SC based on the MICs dataset, **(d)** definition of nodes based on a Schaefer parcellation with higher granularity of 200 nodes, **(e)** using two (*left;* T1-weighted to T2-weighted ratio, principal gradient of functional connectivity) or four (*right;* T1-weighted to T2-weighted ratio, principal gradient of functional connectivity, N-methyl-D-aspartate receptor density, γ-aminobutyric acid type A/Bz receptor density) maps to determine the heterogeneity of regional parameters, and **(f)** including the inter-hemispheric connections in the goodness-of-fit. In all the panels the statistics indicate spatial correlation of each map with the 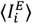 age effect observed using the default configurations at subject level (panel **c** correlated with panel **a**) or age-group level (panels **d-f** correlated with panel **b**).

#### Parcellation, heterogeneity maps and inter-hemispheric connections

Following, we assessed a range of alternative modeling configurations, which we performed at the level of age groups (N = 30, each with 25-27 subjects) to limit the usage of computational resources. We found largely consistent 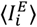 age effect maps similar to the age effects obtained using the main analysis configuration (Figure 5b, S9) with: i) using an alternative parcellation with higher granularity of 200 nodes^60^ (r = 0.479, p_spin_ < 0.001; Figure 5d, S10), ii) using two (r = 0.339, p_spin_ < 0.001) or four (r = 0.832, p_spin_ < 0.001) instead of six heterogeneity maps to determine regional variability of the parameters 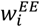 and 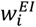 (figure 5e, S11), and iii) including the inter-hemispheric connections in the calculation of goodness-of-fit and the optimizer cost function (r = 0.753, p_spin_ < 0.001; Figure 5f, S12).

#### Conduction velocity

Next, we assessed the potential impact of inter-regional conduction delay at the individual level by repeating the optimal simulations while adding conduction delay between model regions informed by the subject-specific tractograms. We next calculated the node-wise median absolute deviation intraclass correlation coefficient (ICC) of 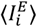 between delayed-conduction and non-delayed-conduction simulations, and found its high test-retest reliability with a mean of 0.975±0.027 using velocity of 1 m/s to 0.998 ±0.003 using velocity of 6 m/s (Figure S13).

#### Gaussian noise random seed

Using a similar analysis as above, we investigated the impact of the array of Gaussian noise injected into the simulations by repeating the optimal simulation of each subject with 50 alternative random seeds for the Gaussian noise. The median node-wise ICC of 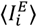 between the original simulations and each of the 50 alternative simulations with different seeds, had an average of 0.802±0.100 (range: [0.449-0.929]) across regions which indicates moderate to high test-retest reliability of 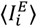 across random seeds (Figure S14).

#### Optimization random seed

It could be that within the parameter space multiple local optima exist that feature different 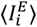 values. To assess whether this may have been the case, we calculated the node-wise ICC of 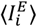 across the two optimal points obtained from the two CMA-ES runs of each subject. We found a mean ICC of 0.947±0.019 (range [0.853-0.984]) across nodes, indicating high test-retest reliability of 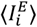 across alternative optima, which shows a low risk of optimizers convergence to different local optima with respect to 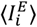 (Figure S15).

### Alternative BNM-based measures of the E-I balance

*In vivo* estimation of the E-I balance based on BNMs has been the aim of several previous studies using this or similar models^47,56,80–82^. Yet, there has been no consensus on the BNM-based measures of the E-I balance and various measures have been proposed and utilized across studies. Here, we present our findings regarding alternative BNM-based measures of the E-I balance used in the previous literature and highlight some considerations regarding their usage.

Firstly, the optimal model parameters have been commonly used as measures of the E-I balance^47,80–82^. Variation of these parameters can be interpreted as shift of the balance towards higher excitation (e.g., increase of *G*or *w*^*EE*^) or higher inhibition (e.g., increase of *w*^*EI*^or *w*^*IE*^). However, in a multidimensional model in which these parameters can simultaneously covary and may be degenerate, the interpretation of their variations is not straight-forward. Indeed, across optimal simulations of subjects in the PNC dataset we found significant associations between optimal parameters, such as a negative association of *w*^*EI*^ and *w*^*IE*^, indicating that lower excitatory-to-inhibitory connection weights are accompanied by (compensatory) higher inhibitory-to-excitatory connection weights (Figure 6a). These associations were also reflected in the effects of age on the parameters (Figure 6b), as for example, there was an inverse correlation between the unthresholded effects of age on *w*^*EI*^ and *w*^*IE*^ (r = -0.667, p_spin_ < 0.001). The observed covariance between model parameters and their age effects, indicates that these age effects should not be interpreted in isolation, and based on this data, the net effect of age on the E-I balance remains ambiguous.

**Figure 6.**
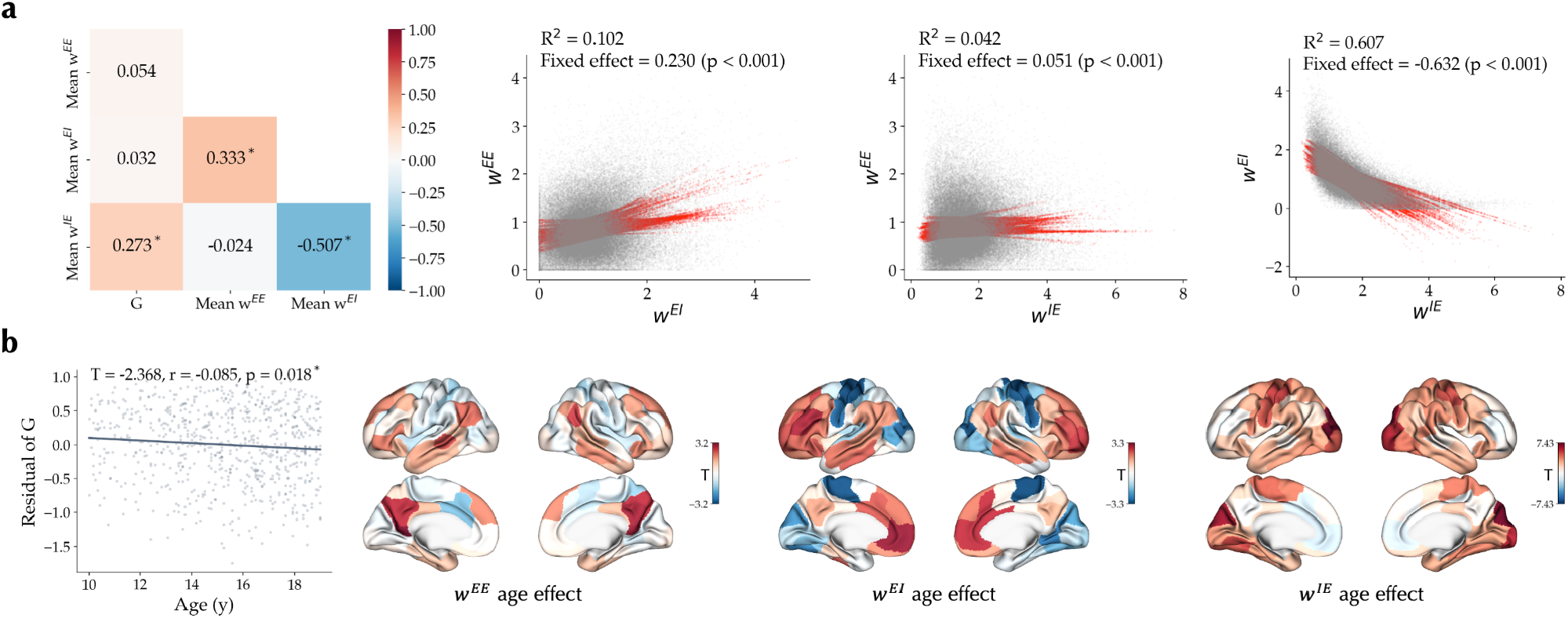
Optimal model parameters inter-relation and association with age in the PNC dataset. **(a)** *left*: Pearson correlation of model parameter *G*and brain-averaged values of regional parameters *w*^*EE*^, *w*^*EI*^ and *w*^*IE*^ across subjects are shown. Asterisks denote statistically significant correlations. *right*: Inter-relation of regional values of parameters *w*^*EE*^, *w*^*EI*^ and *w*^*IE*^ across nodes and subjects based on a linear mixed effects model with random intercepts and slopes per each node. **(b)** *left*: Effect of age on optimal parameter *G*. Points represent the residual of *G*for each subject after removing confounds. *right*: Unthresholded effect of age on regional parameters *w*^*EE*^, *w*^*EI*^ and *w*^*IE*^.

In our study, as a solution to this problem of degeneracy between model parameters, we focused on a state variable of model nodes within the optimal simulations, which, as a “final common pathway”, reflects the collective outcome of the various model parameters on the E-I balance within each node. At a neuronal level, the E-I balance is commonly defined as the balance, or the ratio, between excitatory and inhibitory currents, potentials or conductances onto excitatory neurons^83,84^. Accordingly, in our study, we quantified the E-I balance based on time-averaged input current onto the excitatory neurons, 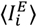, which reflects the net difference of the excitatory and inhibitory currents onto these neurons. Of note, our measure differs from another BNM-based measure of the E-I balance based on model state variables which was utilized in a similar previous study^56^: the ratio of time-average excitatory synaptic gating variable, 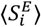, to time-average inhibitory synaptic gating variable,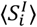. By inspecting the time series of model state variables in an example optimal simulation, we found that 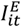 and 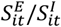 are indeed positively correlated (R^2^ = 0.472). However, in subsequent analyses we found notable differences between these two alternative BNM-based measures of the E-I balance: (i) Assuming that the firing rate of excitatory neurons, *r*^*E*^, is an outcome of the E-I balance^61^ which indicates low versus high states of activity^83^, we expect a measure of the E-I balance to positively correlate with it. Accordingly, in an example simulation and using an exponential generalized linear mixed effects model we found that 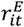 correlates positively across nodes and time with both 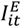 (R^2^ = 0.976) and 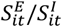 (R^2^ = 0.582), but is more strongly correlated with 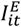 (Figure S16). This was not surprising given model Eq. 3 directly relates 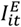 to 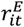. (ii) Next, given the optimal simulation of 40 randomly selected subjects of the PNC dataset, we performed perturbed simulations in which one of the model parameters was increased or decreased by 10%, pushing the simulation to an expected state of increased/decreased excitation/inhibition (e.g., 10% increase of *G*is expected to push the system towards higher excitation). We then used paired T-tests to compare each measure of the E-I balance before and after the perturbation (Figure S17), and found larger effects of perturbations on 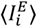 (mean |T| = 21.476±5.375), compared to 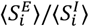 (mean |T| = 6.995±2.730). This shows that 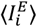 may be more sensitive to capture the variations in the E-I balance caused by parameter perturbations. (iii) Lastly, we studied the effects of age on 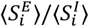 in the PNC dataset across different model configurations and found: 1) widespread and unimodal-dominant increases using main configurations, 2) increases in unimodal and decreases in transmodal areas when using template SCs, and 3) widespread and unimodal-dominant decreases using age-group-averaged functional data, template SC, and two heterogeneity maps, which reflected the pattern observed in the study by Zhang et al.^56^ that used a similar configuration (Figure S18). This showed that 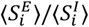 age effects vary with different modeling configurations, though we found it was less sensitive to the effect of random seeds in the simulation (mean ICC: 0.999) and optimization (mean ICC: 0.968) relative to 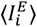.

These findings, combined with the commonly used definition of the E-I balance in experimental and theoretical research at neuronal level, suggest that 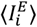 may be a more direct and interpretable measure of the E-I balanced using the BNM approach compared to the alternatives used in the literature. They also highlight how modeling choices and parameters can impact the outcomes of E-I changes with age.

## Discussion

In this study, we used large-scale simulations of biologically realistic and individualized biophysical network models to estimate regional excitation-inhibition balance based on *in vivo* imaging data and evaluated its maturation during adolescence. We found a developmental shift of the E-I balance towards higher inhibition (lower excitation) in the association areas while the sensorimotor areas showed a lack of significant changes or a developmental shift towards higher excitation (lower inhibition). This finding was supported by imaging data from two independent datasets and through investigating both cross-sectional, inter-individual age-related variations of the E-I balance, as well as its longitudinal, within-individual changes through adolescence. Our observed pattern of regional variability in the E-I balance development aligned with the sensorimotor-association axis of cortical organization, and highlighted divergence of low-versus high-level brain functions, as well as early-versus late developmental timing of the sensorimotor and association areas. Lastly, we showed that our findings were robust to various modeling nuisances and choices and contrasted our simulation-based measure of the E-I balance to the alternative measures used in the literature.

The robust and replicable developmental pattern of decreased 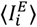 found in the association areas, indicates a relative shift of the E-I balance towards increased inhibition or decreased excitation. This observation is in line with several findings from previous animal and human studies. At the molecular level, transcriptomics and proteomics analyses have revealed peri-adolescence changes in the expression of genes related to excitation and inhibition, such as the NMDA receptor subunits^85^, calcium-binding proteins parvalbumin, calretinin, and calbindin, which are expressed in different types of interneurons^9,26,27,32^, and GABA_A_ receptor subunits^28–31^. These molecular shifts mirror changes in neuronal functional properties. For instance, within the prefrontal cortex, there is an increase in the subunit composition of the GABA_A_ receptors, from *α*_2_- to *α*_1_-containing receptors, which have a faster decay time, resulting in faster synaptic inhibition^31,32,86^. Consistent with this, recording of pyramidal neurons of the prefrontal cortex in non-human primates has indicated an increase in the strength as well as shortening of the inhibitory postsynaptic currents^9,30^. At the same time, microscopic investigation of the pyramidal neurons in the prefrontal cortex across different species has revealed a dramatic reduction of the excitatory synaptic density during adolescence^19–23,25,87^. In humans, magnetic resonance spectroscopy has been used in several studies to quantify *in vivo* levels of glutamate and GABA, primarily in the frontal areas. Yet, the findings of these studies have been inconsistent and include reports of age-related decrease of glutamate^39,88,89^ as well as increase^38,40^ or decrease of GABA^39^, increase of glutamate-GABA balance^39,89^, and several null findings^38–40,89–91^. Furthermore, a recent study on the PNC dataset quantified the *in vivo* E-I ratio by modeling multivariate patterns of FC and assessing their (dis)similarity to the FC of adults receiving alprazolam, a GABAergic agonist, and reported a significant developmental increase of the E-I ratio which was specific to association areas^33^.

In contrast to the widespread relative increase of inhibition in association regions during adolescence, we found relative increase of excitation or no significant age-related changes in the sensorimotor areas. Human cortical maturation is suggested to unfold across a sensorimotor-association axis, with a differential temporal patterning indicating earlier maturation of the sensorimotor areas in contrast to later and more protracted maturation of the association cortices^18^. In line with this, we found that the spatial pattern of differential E-I balance maturation across cortical areas co-aligns with the proposed sensorimotor-association axis of neurodevelopment^18^, and indicated that the genes preferentially expressed in association areas are more prominent in later stages of development. The sensorimotor-association neurodevelopmental variation has been observed in cortical maturation of macrostructural features^92^, intracortical myelination^93,94^, white matter connectivity^95^ and functional organization^18,96,97^, parallel to the maturation of excitation and inhibition^18^. Consistent with the findings in the prefrontal cortex^21,22^, accelerated pruning of excitatory synapses around puberty has been observed in sensorimotor areas as well^87,98,99^, though synaptic pruning in association regions is protracted and peaks later than in sensorimotor areas^18,24^. In addition, the maturation of parvalbumin inhibitory interneurons in association areas is suggested to be more prolonged^18,100^. Given the differences in the neurodevelopmental timing along the sensorimotor-association axis, it may be that the E-I balance matures earlier in the sensorimotor areas before adolescence, and hence, we did not find maturational increase of relative inhibition in these areas during our study age period. In line with our observation, two other studies using human *in vivo* markers of E-I balance reported significant increase of inhibition markers in association areas despite no significant changes in sensorimotor regions^33,40^. Therefore, an intriguing area for the future research will be to investigate the maturation of model-estimated E-I balance in earlier stages of development before adolescence.

Our study extends upon a recent modeling study which found widespread relative increase of inhibition across the cortex, most prominently in the sensorimotor areas, by using BNMs constructed for 29 age groups of the PNC dataset and based on a template SC of an adult sample^56^. In contrast, here we used large-scale simulations to construct individualized BNMs^44,54,55^, which allowed more specific simulation-based mapping of the E-I balance in each individual subject and this also enabled studying changes of the E-I balance longitudinally within the same individual. In addition, individualized BNMs are shown to enhance reliability of model parameters and fingerprinting accuracy of the simulated data^55^. Though our findings within the association areas were in agreement with the study by Zhang et al.^56^, indicating a maturational shift of the E-I balance towards inhibition, they diverged in the sensorimotor areas. We suspect this divergence can be attributed to several differences of the two studies, which, in addition to the usage of group-level versus individualized BNMs, include different simulation-based markers of the E-I balance as well as the methodological details of image processing, modeling and optimization. Importantly, we showed that we can (qualitatively) replicate the findings of Zhang et al.^56^ at the level of age groups when similarly using T1w/T2w and FC G1 as the heterogeneity maps and 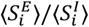 as the marker of E-I balance. However, we presented findings which highlighted that 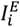 compared to 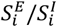, as well as model parameters, may be a more direct BNM-based marker of the E-I balance and be closer to its common definition in the literature as the balance, or the ratio, between excitatory and inhibitory currents, potentials or conductances onto excitatory neurons^83,84^. Indeed, establishing the optimal methodological choices in mapping the E-I balance using BNMs is a challenging open question, and it will be crucial for the future research to: i) comprehensively assess how modeling choices may affect the simulation-derived E-I measures, ii) evaluate the replicability of our findings in alternative datasets, particularly with higher imaging quality and more extensive follow-ups, and the fitting of BNMs to alternative or additional empirical measures, for instance using magneto-/electroencephalography^44,101^, and iii) ultimately, experimentally validate the E-I measures obtained through fitting of BNMs to the fMRI data against empirical measures obtained using electrical recordings, or evaluate them in response to excitatory/inhibitory interventions.

Overall, in this study we observed *in vivo* evidence based on individualized BNMs suggesting a replicable and robust relative increase of cortical inhibition in association areas during adolescence. The normative maturation of E-I balance is suggested to have important functional consequences^18,102^ and its dysmaturation is believed to be associated with various neurodevelopmental disorders^11,12^. For example, the neurodevelopmental model of schizophrenia suggests that aberrant cortical maturation, importantly in the development of excitatory and inhibitory functions, may contribute to the emergence of the disease later in life^12,13,17,103^. Therefore, beyond further methodological and validation studies discussed above, future studies will be needed to investigate the clinical relevance of adolescent maturation of E-I balance in association with the risk and diagnosis of neurodevelopmental disorders such as schizophrenia. Ultimately, investigating model-estimated E-I balance and its deviations from the normative developmental trajectory may be used as a biomarker which can present an opportunity for early detection and treatment of such disorders.

## Methods

This research complies with the ethical regulations as set by The Independent Research Ethics Committee at the Medical Faculty of the Heinrich Heine University Düsseldorf (Study number 2018– 317). We used previously published data from sources that have received ethics approval from their respective institutions^52,53,104,105^.

### Participants

We studied adolescents from two population-based datasets, including the cross-sectional Philadelphia Neurodevelopmental Cohort (PNC)^52,104,105^ and the longitudinal IMAGEN dataset^53^. We selected subjects and follow-up sessions within the age range of 10 to 19 years. In the IMAGEN cohort, within this age range, phenotypic assessments were conducted at the ages of 14, 16 and 19 and imaging data was acquired at the ages of 14 and 19. Next, we excluded subjects with poor quality of raw or processed imaging data, as detailed below in the section “Image processing and quality control”. In the IMAGEN, the subjects were excluded if the imaging data had poor quality in any of the baseline or follow-up sessions, except for the diffusion weighted imaging data for which subjects were selected based on the image quality of the follow-up session. Our final sample consisted of 752 adolescents from the PNC (409 female, mean age: 15.3±2.4 years) and 149 participants from IMAGEN (72 female, baseline mean age: 14.4±0.4 years, follow-up mean age: 18.9±0.5 years). The PNC data was collected at a single center in Philadelphia, while the IMAGEN data was acquired in five different centers across Europe, in Dresden (N = 58), Paris (N = 45), Mannheim (N = 30), London (N = 14) and Dublin (N = 2). Of note, in some of the robustness analyses performed on the PNC we used aggregate data of 30 age groups, which were defined as 29 groups of 25 subjects and one group of 27 subjects, after sorting the subjects by age.

### Image acquisition

T1-weighted, resting-state functional magnetic resonance imaging (rs-fMRI) and diffusion weighted imaging (DWI) data was acquired using 3T scanners from different manufacturers (PNC: Siemens, IMAGEN: Siemens, Philips, General Electric). For more details on the image acquisition parameters we refer the reader to the respective publications for each dataset (PNC: Table 1 of Ref.^52^, IMAGEN: Supplementary Table 5 of Ref.^53^). Of particular relevance to our study, the repetition and acquisition times of the rs-fMRI images were respectively 3 seconds and 6:18 minutes in the PNC, and 2.2 seconds and 6:58 minutes in the IMAGEN.

### Image processing and quality control

#### T1-weighted structural magnetic resonance imaging

T1-weighted MRI images were processed using the *recon-all* command of FreeSurfer (version 7.1.1; https://surfer.nmr.mgh.harvard.edu/), which includes brain extraction, registration to standard space, tissue segmentation, and cortical surface reconstruction^106,107^. Quality of the FreeSurfer output was controlled based on the Euler characteristic, which represents the number of cortical surface defects before correction. We excluded outlier subjects with Euler characteristic greater than Q3 + 1.5 × *IQR* of their cohort^108^. The FreeSurfer output and raw T1w images were subsequently used in the pipelines of rs-fMRI and DWI processing.

#### Resting-state functional magnetic resonance imaging

Resting-state fMRI images were preprocessed using fMRIprep (version 22.0.0; https://fmriprep.org/en/stable/), which performed brain extraction, image registration and motion correction, estimation of confounds, and when possible, susceptibility distortion and slice time corrections^109^. The latter two steps were omitted in the IMAGEN data given unavailability of slice timing and variability of fieldmap formats across centers. We next further processed the output of fMRIprep, which involved the following steps: 1) applying Schaefer-100 parcellation^60^ by taking the average signal of all vertices within each parcel at each time point, 2) removing the first 3 volumes, 3) high-pass temporal filtering > 0.013 Hz, 4) regressing out confounds including average signal of white matter and cerebrospinal fluid voxels as well as 24 motion parameters (translation and rotation in the three directions, in addition to their squares, derivatives, and squares of derivatives)^110^ and 5) scrubbing motion outliers, defined based on root mean squared translation > 0.25 mm. The scrubbing was done by setting the signal in motion outlier volumes to zero while Z-scoring the rest of the volumes. This approach, compared to discarding the motion outliers, preserves the temporal structure of the BOLD signal which is important in calculating dynamic functional connectivity.

We excluded subjects/sessions with high rs-fMRI in-scanner motion defined as less than 4 minutes of scan remaining after scrubbing motion outlier volumes or a time-averaged root mean square > 0.2 mm. In addition, we performed visual quality control of the fMRIprep output and excluded subjects with gross misregistration or incomplete field of view.

#### Functional connectivity

Functional connectivity (FC) was calculated as the Pearson correlation of BOLD signal time series between the cortical areas. In the PNC, we additionally calculated age-group FCs by taking the mean of Z-transformed FC matrices across the subjects of each age group followed by its inverse Z-transformation.

#### Functional connectivity dynamics

Functional connectivity dynamics (FCD) matrix represents the temporal variability of the dynamic patterns of FC computed across sliding windows of time^111^. To compute the FCD matrix, we initially calculated time-resolved FC matrices of sliding windows (PNC: size 30 s and step 6 s, IMAGEN: size 30.8 s and step 4.4 s). We discarded edge windows and windows with ≥ 50% motion outliers. Subsequently, we computed FCD as the correlation between lower triangular parts of window FC patterns. The distribution of values within the FCD matrix represents the amount of recurrence of time-resolved FC patterns. In the PNC, we additionally calculated age-group FCDs by concatenating lower triangles of FCD matrices of all subjects within an age group, followed by its sorting and downsampling to reduce computational costs of model fitting.

#### Diffusion-weighted imaging

DWI images were processed using Micapipe (version 0.1.1; https://micapipe.readthedocs.io/en/latest/)^112^ which combines tools from FSL (version 6.0.0)113 and MRTrix3 (version 3.0.0)114. This pipeline performed DWI processing steps including rigid-body alignment of images, MP-PCA denoising, Gibbs ringing correction, motion and eddy current-induced distortions correction, non uniformity bias correction, registration to the processed structural image, brain mask generation and estimation of fiber orientation distributions using spherical deconvolution. Following, on each image probabilistic tractography was performed using iFOD2 algorithm to estimate 10 million streamlines^115^. Additionally, a track density image was computed using the iFOD1 algorithm with 1 million streamlines which was used for the quality control^116^. The quality control of the tractograms was done by visual inspection of the tractogram density images.

#### Structural connectivity

The structural connectome (SC) matrix for each subject was created using Micapipe by parcellating the tractogram using the Schaefer-100 parcellation map^60^ non-linearly registered to the DWI space. We subsequently normalized each SC matrix by division by its mean × 100, resulting in an equal mean of 0.01 in all SCs.

In addition to subject- and session-specific SCs of the adolescents, we used a higher-quality DWI data (3T, three shells, 140 directions) of an adult sample of 50 healthy volunteers (MICs dataset, 23 female, mean age: 29.5±5.6 years) to construct a template SC^117^. To do so, we obtained the SC of individual MICs subjects preprocessed using Micapipe and calculated a group-averaged SC by taking the median of streamline counts in each edge. The template SC was subsequently normalized by its mean × 100, similar to the subject- and session-specific SCs of the adolescents.

In the individualized models reported in the main analyses, we used subject-specific SCs, but the adult template SC was used in the robustness analyses on the confounding effects of inter-individual variability in SC, as well as the effects of modeling choices. Of note, in IMAGEN, given the lower quality of tractograms in the baseline session, the main analyses were performed by using the follow-up SC of each subject for the modeling of functional data in both the baseline and follow-up sessions. However, in a supplementary analysis, we additionally performed simulations using session-specific SCs, within a subset of the IMAGEN subjects with adequate quality of tractograms in both sessions (N = 110, 52 female).

### Biophysical network modeling

Next, we performed BNM simulation-optimization at the level of each individual subject/session (Figure 1).

#### Model simulation

We simulated the spontaneous neuronal activity of 100 cortical regions from the Schaefer atlas^60^ as network nodes regulated by the reduced Wong-Wang model and interconnected through the SC^61^. In short, this model describes the activity of large ensembles of interconnected excitatory and inhibitory spiking neurons in each area by a dynamic mean field model as a reduced set of dynamic equations governing the activity of coupled excitatory (E) and inhibitory (I) pools of neurons. In this reduced model, the excitatory synaptic currents are mediated by the N-methyl-D-aspartate (NMDA) receptors and the inhibitory synaptic currents are mediated by the γ-aminobutyric acid type A (GABA_A_) receptors. Within each cortical region, the E and I neuronal pools are interconnected, and between regions, the E neuronal pools are coupled through a scaled SC matrix.

##### Model equations

The model is mathematically described by a set of dynamic equations^61^. The total input current (in nA) to each excitatory (E) and inhibitory (I) neuronal pool of each cortical node i, 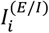, is calculated as:

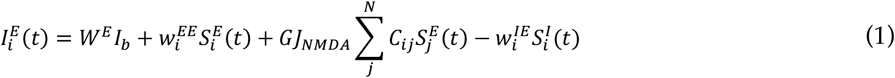

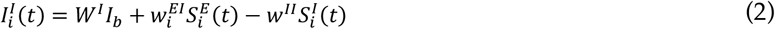

where *W*^*E*^*I*_*b*_ = 0.382 nA and *W*^*I*^*I*_*b*_ = 0.267 nA are the baseline input currents; 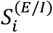 denote the synaptic gating variables; *N* = 100 is the number of nodes; *c* is the structural connectivity matrix, which together with the NMDA receptor conductance, *J*_*NMDA*_ = 0.15 nA, and *G*(global coupling), a free parameter of the model, determine the excitatory input current transmitted from the other nodes;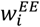 is the recurrent excitatory connection weight; 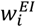 indicates the excitatory-to-inhibitory connection weight; 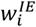 is the inhibitory-to-excitatory connection weight; *w*^*II*^ = 1.0 denotes recurrent inhibitory connection weight. The local connection weights 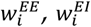 and 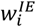 can vary across nodes and simulations through the free parameters of the model and the feedback inhibition control, as described below.

The total input current received by each neuronal pool is subsequently transferred to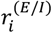, firing rates in Hz, using the sigmoidal neural response function, *H*^(*E*/*I*)^:

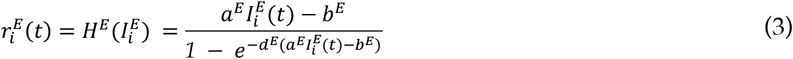

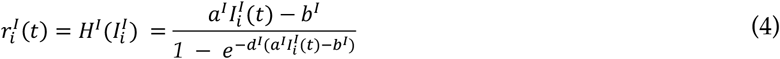

where *a*^*E*^ = 310 nC^-1^ and *a*^*I*^ = 615 nC^-1^ determine the slope of *H*^(*E*/*I*)^; *b*^*E*^ = 0.403 nA and *b*^*I*^ = 0.288 nA define the thresholds above which the firing rates increase linearly with the the input currents; *d*^*E*^ = 0.16 and *d*^*I*^ = 0.087 determine the shape of *H*^(*E*/*I*)^ curvature around *b*^(*E*/*I*)^.

Finally, the synaptic gating variables, 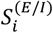, follow:

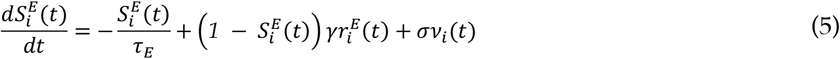

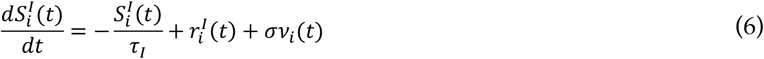

where 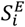 is mediated by NMDA receptors with a decay time constant *τ*_*E*_ = 0.1 s and *γ* = 0.641, and 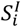 is mediated by GABA receptors with a decay time constant *τ*_*I*_ = 0.01 s. *v*_*i*_ (*t*) is uncorrelated standard Gaussian noise with an amplitude of *σ* = 0.01 nA. 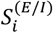 is subsequently bound within the range [0, 1]. Note that the fixed parameters in equations 1-6 are based on a previous paper by Deco et al.^61^.

##### Model free parameters

The model is controlled by 15 free parameters, including *G*, as well as bias terms and coefficients for a fixed set of six biological maps which together determine the regional values of 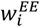 and 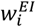. More specifically, 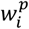, *p* ∈ {*EE, EI*} is calculated as:

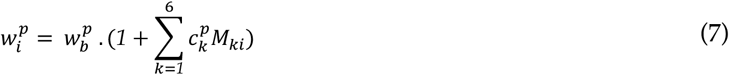

where 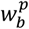, the bias term, and *c*^*p*^, a vector of six coefficients, are free parameters. *M* is a 6×100 matrix including the Z-scored biological maps in Schaefer-100 parcellation (Figure S1). These maps were based on independent samples of healthy individuals, obtained from *neuromaps*^74^ and Hansen et al.^70^ and included: 1) group-averaged T1-weighted/T2-weighted (T1w/T2w) ratio map of the Human Connectome Project (HCP) dataset^68,69,118^, 2) group-averaged cortical thickness map of the HCP dataset, 3) principal gradient of functional connectivity (FC G1)^72^, 4) principal axis of Allen Human Brain Atlas gene expression data (Gene PC1)^71,73^, 5) NMDA receptor density positron emission tomography map^67,70^ and 6) GABA_A/BZ_ receptor density positron emission tomography map^70,75^. The resulting 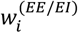 maps were subsequently shifted if needed to ensure 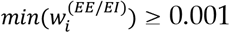. We used the following ranges for the model free parameters: *G*: [0.5, 4.0], 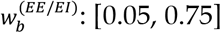. The range of coefficients for each map was defined as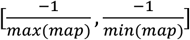, corresponding to c_T1w/T2w_: [-0.48, 0.59], c_cortical thickness_: [-0.40, 0.39], c_FC G1_: [-0.59, 0.72], c_Gene PC1_: [-0.36, 0.48], c_NMDA_: [-0.49, 0.42] and c_GABAa/bz_: [-0.30, 0.32].

##### Feedback inhibition control

Feedback inhibition control (FIC) was used to determine the regional values of 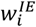 in each simulation, given the SC and other model parameters. The FIC algorithm aims to maintain a state of E-I balance in each region by adjusting 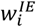 to satisfy an excitatory firing rate close to 3 Hz, which is suggested to be within the biological range^61^. We used a two-stage implementation of the FIC by combining the original numerical implementation^61^ with an analytical solution proposed by Demirtaş et al.^66^. The latter solution analytically solves for 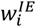 to satisfy the self-consistency of the model equations at the steady-state condition with 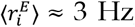, corresponding to 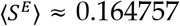and 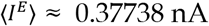:

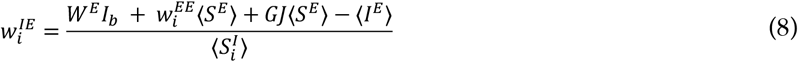

in which the steady-state inhibitory synaptic gating variable 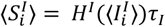 was estimated by solving for 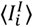 in:

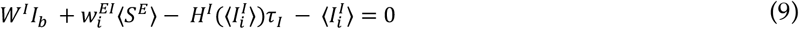

Subsequently, analytical estimates of 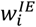 values were fed into the numerical implementation of FIC and were adjusted numerically^61^. In this approach, given the analytical estimates of 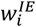, the model equations 1-6 are numerically integrated for a short period of 10 s and subsequently the average input current to the excitatory pool of each brain region, 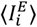 is calculated. If 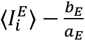 in a region exceeded the target - 0.026 by more than 0.005 nA, 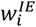 is up-/downregulated when the input current is higher/lower than the target, and the simulation is repeated with the adjusted 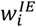 values in the next trial. This procedure is repeated for 10 trials or until the FIC target is satisfied in all nodes. Note that the maximum number of FIC numerical adjustment trials used here is lower than that of the original implementation to facilitate the scaling of the simulations. Furthermore, as the initial 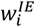 values are estimated analytically rather than being fixed to 1 (as was done in the original implementation), a smaller number of trials will be needed.

#### Hemodynamics model

The simulated synaptic activity of the excitatory population of each node, 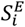, was subsequently fed to the Balloon-Windkessel model of hemodynamics to simulate BOLD signal^119^. This model is mathematically described by the following system of differential equations with state variables *x* (vasodilatory signal), *f* (blood inflow), *v* (blood volume) and *q* (deoxyhemoglobin content):

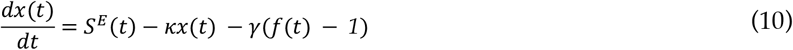

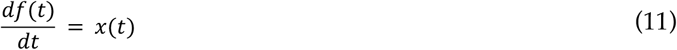

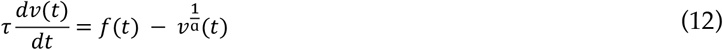

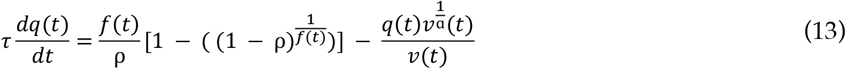

where 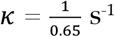 is rate of signal decay, 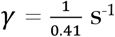 is rate of flow-dependent elimination, *τ* = 0.98 s is hemodynamic transmit time, *α* = 0.32 is Grubb’s exponent and ρ = 0.34 is resting oxygen extraction fraction. These parameters were based on a previous paper by Friston et al.^119^. Finally, based on the model state variables the BOLD signal is calculated as:

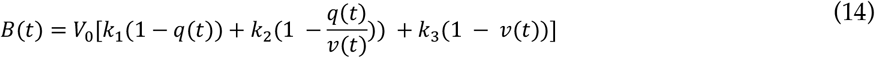

in which *V*_0_ = 2% is the resting blood volume fraction^119^ and *k*_1_ = 3.72, *k*_2_ = 0.527 and *k*_3_ = 0.53 are dimensionless parameters that were derived for 3T scans^66,120^. Last, the simulated BOLD signal was downsampled to match the repetition time of the empirical rs-fMRI data, that is, 3 s for the PNC and 2.2 s for IMAGEN.

#### Numerical integration of the models

For each simulation, the model equations were numerically integrated using the Euler method with a time step of 0.1 ms for the neuronal model (Equations 1-6) and a time step of 1 ms for the hemodynamic model (Equations 10-13). The model simulations were performed using in-house code (https://github.com/amnsbr/bnm_cuda, https://cubnm.readthedocs.io) on graphical processing units (GPUs) of JURECA-DC, a supercomputer at the Jülich Supercomputing Centre^121^. This GPU implementation enabled efficient parallelization of calculations for individual simulations (across GPU ‘blocks’) and the regions within each simulation (across GPU ‘threads’). To match the duration of empirical rs-fMRI scans, the simulations were done for a biological duration of 450 s, from which the first 30 s were discarded to ensure the BNM system’s state has stabilized.

##### Model evaluation

The goodness-of-fit of the simulated BOLD signal given a set of candidate parameters and SC matrix to a target empirical BOLD signal was evaluated based on three measures of static and dynamic functional connectivity, following previous studies^56,65^:

###### Static edge-level functional connectivity

The simulated FC was calculated as the correlation of simulated BOLD signal time series between nodes. The correspondence of simulated and empirical FC patterns was evaluated by calculating the Pearson correlation coefficient between the lower triangles of the matrices (FC_corr_), with larger values representing higher correspondence.

###### Static global functional connectivity

The absolute difference of the averaged simulated and empirical FC matrices across all the lower triangular edges (FC_diff_) was calculated to assess the similarity of global FC strength, with smaller values showing higher correspondence.

###### Functional connectivity dynamics

The simulated FCD matrix was constructed by calculating the correlation of FC patterns between sliding windows of simulated BOLD signals, as described previously for the empirical data. The correspondence of simulated and empirical FCD distributions was calculated as the Kolmogorov-Smirnov (KS) distance of their lower triangular parts (FCD_KS_), with smaller values showing higher similarity of the distributions.

Subsequently, these measures were combined into a single measure of goodness-of-fit as FC_corr_ – FC_diff_ - FCD_KS_. Of note, in goodness-of-fit calculations, following Demirtaş et al.^66^, we excluded the interhemispheric connections. However, we also performed a robustness analysis in which these connections were included in the goodness-of-fit calculations.

##### Parameter optimization

The model free parameters (N = 15) were fit to the empirical data of each subject/session using the covariance matrix adaptation-evolution strategy (CMA-ES) optimization algorithm^62–64^. CMA-ES is an efficient evolutionary optimization algorithm which features a set of Λ particles exploring the parameter space collaboratively in an iterative process. The particles from each iteration, which are individual simulations with different free parameters, are regarded as a generation from which only the best particles are selected to form the descendant population of the next generation. Specifically, at each generation the cost function of each particle is calculated, as described below. Following, a weighted mean of the best fitting ⌊Λ/2⌋ particles is calculated. Then, a new generation of particles is created by taking Λ samples from a multivariate normal distribution centered around the weighted mean of the best fitting ⌊Λ/2⌋ particles from the previous generation. The covariance is determined by a matrix which is updated to take the location of the current best points into account. In this way, the search distribution is adapted iteratively towards a concentration around the optimal solutions. This iterative process is continued for a maximum of 80 generations, following Wischnewski et al.^64^, and eventually, the optimal point across all generations is selected as the optimal parameters for the best fit of the simulation to the given target empirical data. We also applied an early termination rule in which the iteration was stopped if there was no improvement in the cost function greater than 0.005 over the past 30 generations.

The optimization goal was to maximize the goodness-of-fit while minimizing a penalty term. Particles were penalized if 1) the parameter of sampled particles fell outside the pre-specified ranges, in which case the parameters were corrected and a penalty was added^63^, or 2) the 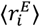 was outside the range of 2-4 Hz in any node, indicating insufficiency of the FIC. For the latter, the FIC penalty was calculated as:

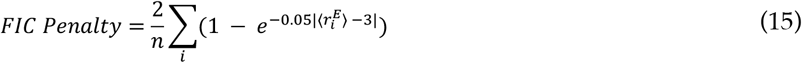

in which *n* is the number of nodes and the summation is done across nodes with out-of-range 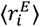. Of note, we refrained from setting the success of FIC as a hard constraint, but through this penalty term applied it as a soft constraint, to allow for inter-individual variability of E-I balance while keeping it within a biologically viable range^61^.

Given the relatively high dimensionality of the optimization problem in our model and to sufficiently cover the large parameter space, we chose Λ = 210 in the CMA-ES. Consequently, with maximum 80 generations, this involved performing a maximum of 16800 simulations per run for each subject/session, which necessitated an efficient GPU-based implementation. In addition, for each given subject/session, the model simulation-optimization was repeated twice using different random seeds of the optimization to assess and reduce the risk of local optima. We then took the best fitting simulation across the two runs of each subject/session as their optimal simulation for the next step.

##### Estimation of the excitation-inhibition balance *in silico*

Thus far, we described the procedure for deriving the optimal parameters that result in a simulation best fitting to the empirical rs-fMRI data of a given subject/session using individualized BNMs. Following, given the optimal simulation for each subject/session, we extracted an *in silico* measure of regional E-I balance. To do so, we calculated the average of total input current to the excitatory neurons of each region, after discarding the initial 30 seconds of the simulation. This measure can be interpreted as an *in silico* marker of the regional E-I balance, as 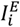 (Eq. 1) results from the combination of excitatory input currents to each node (from itself, and from the excitatory neurons of the other nodes through the SC) balanced by the inhibitory currents from the inhibitory neuron of the same node. Therefore, an increase of 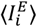 can be interpreted as a relative increase of excitation or decrease of inhibition.

Furthermore, using a similar approach we calculated 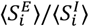 as an alternative marker of the E-I balance used in a previous study^56^.

##### Perturbed simulations

We performed a control analysis to assess the effect of known perturbations in the model parameters on the alternative markers of the E-I balance, namely, 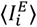 and 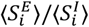. In this analysis, we randomly selected 40 subjects of the PNC dataset and for each subject, given their optimal simulations, we performed perturbed simulations in which one of the model parameters (*G, w*^*EE*^, *w*^*EI*^ or *w*^*IE*^) was increased or decreased by 10% while the other three parameters were fixed to the optimal values. Notably, in these simulations, when *w*^*IE*^ was not perturbed, it was not readjusted using FIC, and was fixed to the *w*^*IE*^ values obtained by the FIC run on the original optimal simulation. Similarly, perturbation of *w*^*IE*^ was done by a 10% increase or decrease of these original *w*^*IE*^ values. This was to ensure only one parameter in the model is perturbed and therefore the net direction of the effect of perturbation on the E-I balance is predictable. Lastly, the effect of perturbation on the E-I balance markers in each subject was assessed using paired T-tests comparing the marker across nodes before and after the perturbation.

### Statistical and contextualization analyses

#### Age effects

Given the regional *in silico* regional measures of the E-I balance for each subject/session, 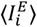, we performed group-level univariate statistical analyses to investigate the effects of age on these measures. Linear regression models were used to study the effect of age on 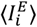 and map coefficients, with goodness-of-fit, sex, and rs-fMRI in-scanner motion (based on time-averaged root mean squared translation) as confounds. In IMAGEN, longitudinal variation of E-I measures across the two sessions was assessed using similar mixed effects linear regression models with random intercepts per subject and inclusion of site as an additional confound. In each model we excluded outliers with a dependent variable ≥3 SDs above/below the mean. We adjusted for multiple comparisons across regions using false discovery rate (FDR) based on the Benjamini/Hochberg method (q < 0.05). Similar models were used to investigate the effects of age on optimal model parameters and 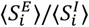. We used *statsmodels*^122^ (https://www.statsmodels.org/stable/index.html) to perform regressions and FDR adjustment.

#### Within-sample stability of age effects using subsampling

In each dataset, we randomly selected 100 subsamples of the subjects (N within each subsample: 376 in PNC, 74 in IMAGEN) and investigated the age effects of 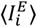 separately in each subsample. Subsequently, we calculated the correlation of unthresholded age effect maps between all pairs of subsamples, and reported its distribution as a measure of within-sample stability.

#### Spatial correlation of maps

We calculated spatial correlation of maps using spin permutation in i) assessing between-sample replicability of PNC and IMAGEN 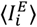 age effect maps, ii) assessing stability of 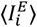 age effect maps within the PNC in the different robustness analyses in comparison to the age effect map observed in the main analysis, iii) assessing the correlation of 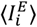 age effect maps with the sensorimotor-association axis of cortical organization described in a recent study by Sydnor et al.^18^ as well as its components or their substitutes^68,69,71–73,77,123–128^ (Figure S6, Table S1), and iv) NeuroSynth^77^ meta-analytical maps of 123 cognitive and behavioral terms (Table S2, based on Hansen et al.^70^) obtained from BrainStat^78^. The target maps in the latter two analyses were available in *fsaverage, fsLR*, or MNI152 spaces and were parcellated in their respective spaces using the Schaefer-100 parcellation^60^. Spin permutation was implemented at parcel level using the ENIGMA Toolbox^129^ (https://github.com/MICA-MNI/ENIGMA) and using 1000 permutations.

#### Distribution of the 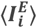 age effects across the canonical resting-state networks

We assessed the association of the 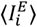 age effect maps with the map of seven canonical resting-state networks^76^ using one-way ANOVA with post-hoc Bonferroni-corrected independent T-tests. To control for spatial autocorrelation we assessed the statistical significance of resulting F and T statistics using null distributions generated from 1000 spun surrogates based on parcel-level spinning implemented in the ENIGMA Toolbox^129^.

#### Partial least squares regression of gene expressions and their developmental enrichment

Regional microarray expression data were obtained from 6 post-mortem brains (1 female, age = 24.0– 57.0) provided by the Allen Human Brain Atlas^71^ (AHBA; https://human.brain-map.org). Data was processed with the *abagen* toolbox^73^ (https://abagen.readthedocs.io/en/stable/) using the Schaefer-100 atlas^60^. Gene expression data from the right hemisphere was excluded due to the sparsity of samples and large number of regions with no expression data. We subsequently used *scikit-learn*^130^ (https://scikit-learn.org/stable/) and performed partial least squares regression analysis to identify gene expression patterns with high spatial co-alignment with the 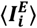 age effect maps within the left hemisphere. After selecting the top 500 genes with highest absolute weights, we divided them into two sets of positively- and negatively-associated genes. Subsequently, using an online tool we performed specific expression analysis across developmental stages and brain structures^79^ (http://doughertylab.wustl.edu/csea-tool-2/). This tool performs comparison of the gene lists against developmental expression profiles from the BrainSpan Atlas of the Developing Human Brain (http://www.brainspan.org), and for each developmental stage and brain structure reports the inverse log of Fisher’s exact p-values. Here, for each set of genes we reported the inverse log of p-values based on the specificity index (pSI) threshold of 0.05 within the cerebral cortex.

#### Test-retest reliability of 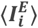 across simulations

We measured node-level test-retest reliability of 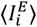 across simulations of the same subject performed using i) alternative optimizer seeds, ii) alternative Gaussian noise seeds, and iii) alternative conduction delays. This was done by measuring median absolute deviation intraclass correlation coefficient (ICC) of each region between the alternative simulations and across subjects (https://warwick.ac.uk/fac/sci/statistics/staff/academic-research/nichols/scripts/matlab/madicc.m).

#### Association of optimal model parameters across subjects

The association of optimal regional parameters (*w*^*EE*^, *w*^*EI*^and *w*^*IE*^) across subjects and nodes was tested via mixed effects linear regressions with random intercepts and slopes per each node. These regressions were performed using *lme4* R package^131^. In addition, we used Pearson correlation to test the association of the optimal *G*with the mean of optimal regional parameters.

#### Association of model state variables

We randomly selected one subject from the PNC dataset and evaluated the association of state variables within its optimal simulation across time points and nodes. The model state variables 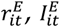 and 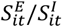 were sampled every TR after the initial 30 seconds of the simulation was removed. Subsequently, as 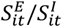 approximates infinity when 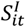 is close to zero, we excluded the data points at the top 2.5 percentile of 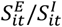. Subsequently we used mixed effects models with or without a logarithmic linking function to test the linear or exponential associations of 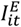 with 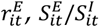 with 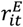 and 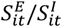 with 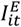, and reported the results of the model with lower Akaike information criterion. The mixed effects models included random intercepts and slopes and were implemented using *lme4* R package^131^ and its Python interface in *pymer4* package.

### Robustness analyses

We performed the following robustness analyses on the PNC dataset:

#### Inter-individual variability of structural connectome

We assessed the potential effect of inter-individual variability of SCs in our findings by performing the following analyses: i) We studied the age effect of 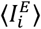 which was additionally controlled for the SC strength of each node, calculated as the row-wise sum of the SC. ii) We performed BNM simulation-optimization using subject-specific functional data as the target, but with the template SC of MICs dataset determining the connectivity of model nodes, thereby eliminating the inter-individual variability of SCs as a potential source of variability in 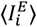. This higher-quality DWI data from an adult sample was chosen to additionally assess the robustness of our results to potential inaccuracies of subject-level SCs derived from relatively lower quality DWI data of the adolescent datasets.

#### Parcellation, heterogeneity maps and inter-hemispheric connections

In these robustness analyses, we performed the BNM simulation-optimization using alternative modeling choices, including: i) the Schaefer-200 parcellation, ii) using T1w/T2w and FC G1 as the heterogeneity maps, iii) using T1w/T2w, FC G1, NMDA and GABA_A/BZ_ as the heterogeneity maps, and iv) including inter-hemispheric connections in the goodness-of-fit and cost calculations. To reduce the computational costs, these analyses were performed at the level of age groups by using age-group functional data and the template SC determining the connectivity of model nodes. Subsequently, we compared the goodness-of-fit measures as well as the 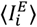 age effect maps derived from these alternative models to the output of BNM simulation-optimization done with the default configurations on the same age-group-level data.

#### Effects of Gaussian noise seed and inter-regional conduction delay

These analyses were performed at the level of individual subjects, by repeating the optimal simulation of each subject obtained in the main analyses with alternative configurations, including: i) using 50 different randomization seeds for generating the Gaussian noise injected into the model (Eq. 5, 6), and ii) adding conduction delay in the signal transmission between the model nodes. For the latter, delay was calculated as SC edge length obtained from the tractography of each subject, divided by a conduction velocity. For each subject we performed six alternative simulations with variable conduction velocities in the range of {1, …, 6} m/s. Of note, in simulations with conduction delay a recent history of 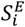 in all nodes needs to be stored in memory, so that the input of node *j* to node *i* at time *t* can be determined based on 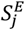 at *delay*_*ij*_ time points ago. Accordingly, to reduce memory needed for storing this history, we performed these simulations by updating the global input to each node at intervals of 1 millisecond, instead of 0.1 milliseconds used in the main analyses. Subsequently, we calculated the goodness-of-fit measures of the alternative simulations to the empirical data of each subject and compared them with the goodness-of-fit measures of the main simulation. In addition, we calculated the ICC of 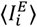 between the main simulation and each of the alternative simulations. In the case of Gaussian noise seeds, we combined the goodness-of-fit as well as ICC measures of the 50 different seeds by taking their median.

## Data and code availability

All the code used in this study for image processing, biophysical network modeling and the statistical analyses is available in a GitHub repository (https://github.com/amnsbr/eidev). Notably, while the main biophysical network modeling in this study were performed using https://github.com/amnsbr/bnm_cuda, we recommend and have included scripts to run the simulation-optimization using the cuBNM toolbox (https://github.com/amnsbr/cuBNM, https://cubnm.readthedocs.io), which we developed using a similar core code but with an improved interface. The adolescent datasets used in this study are restricted-access and cannot be shared openly. The neuroimaging and phenotypic data of Philadelphia Neurodevelopmental Cohort is available as part of the “Neurodevelopmental Genomics: Trajectories of Complex Phenotypes” study (dbGaP Study Accession: phs000607.v3.p2). Our access to this dataset was granted under accession number 32685. The IMAGEN dataset is similarly available to researchers by submitting a data access application to https://imagen-project.org/the-imagen-dataset/. The MICs dataset is openly available at https://osf.io/j532r/.

## Acknowledgements

AS and SLV were funded by the Max Planck Society (Otto Hahn award) and Helmholtz Association’s Initiative and Networking Fund under the Helmholtz International Lab grant agreement InterLabs-0015, and the Canada First Research Excellence Fund (CFREF Competition 2, 2015–2016) awarded to the Healthy Brains, Healthy Lives initiative at McGill University, through the Helmholtz International BigBrain Analytics and Learning Laboratory (HIBALL). LDL was supported by the Federal Ministry of Education and Research (BMBF) and the Max Planck Society (MPG), Germany. SBE was supported by the Deutsche Forschungsgemeinschaft (DFG, EI 816/21-1), the National Institute of Mental Health (R01-MH074457), and the European Union’s Horizon 2020 Research and Innovation Programme under Grant Agreement No. 945539 (HBP SGA3).

The authors gratefully acknowledge computing time on the supercomputer JURECA^121^ at Forschungszentrum Jülich under grant no. “eidev”. This research has been conducted using data from the dbGaP Study Accession: phs000607.v3.p2. Support for the collection of the data for Philadelphia Neurodevelopment Cohort (PNC) was provided by grant RC2MH089983 awarded to Raquel Gur and RC2MH089924 awarded to Hakon Hakonarson. Subjects were recruited and genotyped through the Center for Applied Genomics (CAG) at The Children’s Hospital in Philadelphia (CHOP). Phenotypic data collection occurred at the CAG/CHOP and at the Brain Behavior Laboratory, University of Pennsylvania. This study additionally used the IMAGEN dataset, which received support from the following sources: the European Union-funded FP6 Integrated Project IMAGEN (Reinforcement-related behaviour in normal brain function and psychopathology) (LSHM-CT-2007-037286), the Horizon 2020 funded ERC Advanced Grant ‘STRATIFY’ (Brain network based stratification of reinforcement-related disorders) (695313), Horizon Europe ‘environMENTAL’, grant no: 101057429, UK Research and Innovation (UKRI) Horizon Europe funding guarantee (10041392 and 10038599), Human Brain Project (HBP SGA 2, 785907, and HBP SGA 3, 945539), the Chinese government via the Ministry of Science and Technology (MOST). The German Center for Mental Health (DZPG), the Bundesministerium für Bildung und Forschung (BMBF grants 01GS08152; 01EV0711; Forschungsnetz AERIAL 01EE1406A, 01EE1406B; Forschungsnetz IMAC-Mind 01GL1745B), the Deutsche Forschungsgemeinschaft (DFG grants SM 80/7-2, SFB 940, TRR 265, NE 1383/14-1), the Medical Research Foundation and Medical Research Council (grants MR/R00465X/1 and MR/S020306/1), the National Institutes of Health (NIH) funded ENIGMA-grants 5U54EB020403-05, 1R56AG058854-01 and U54 EB020403 as well as NIH R01DA049238, the National Institutes of Health, Science Foundation Ireland (16/ERCD/3797). NSFC grant 82150710554. Further support was provided by grants from: - the ANR (ANR-12-SAMA-0004, AAPG2019 - GeBra), the Eranet Neuron (AF12-NEUR0008-01 - WM2NA; and ANR-18-NEUR00002-01 - ADORe), the Fondation de France (00081242), the Fondation pour la Recherche Médicale (DPA20140629802), the Mission Interministérielle de Lutte-contre-les-Drogues-et-les-Conduites-Addictives (MILDECA), the Assistance-Publique-Hôpitaux-de-Paris and INSERM (interface grant), Paris Sud University IDEX 2012, the Fondation de l’Avenir (grant AP-RM-17-013), the Fédération pour la Recherche sur le Cerveau.

