## Supplemental Information for "Adolescent maturation of cortical excitation-inhibition balance based on individualized biophysical network modeling"

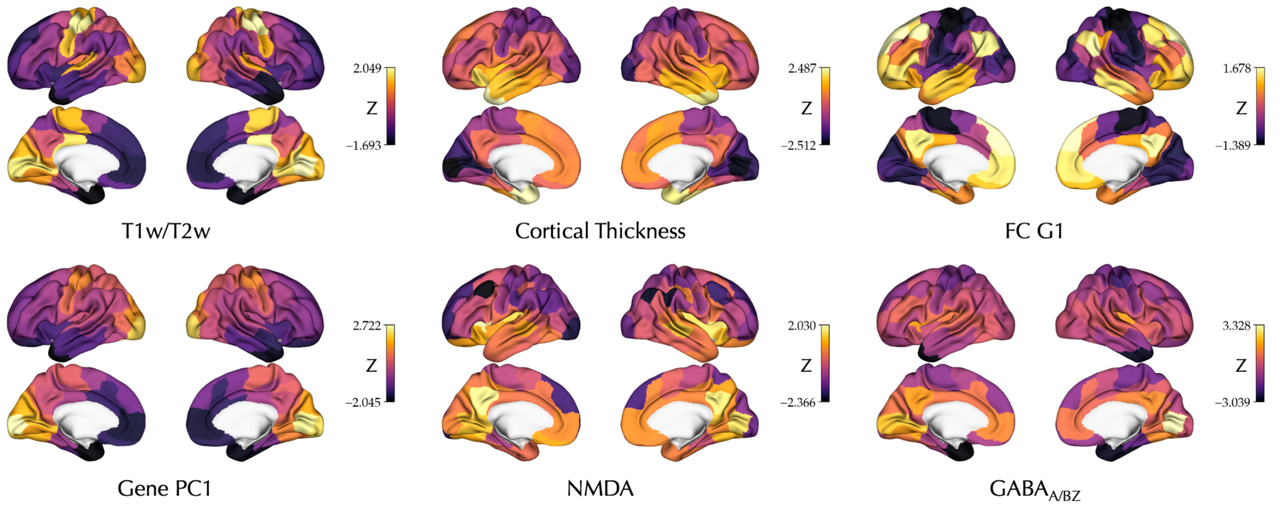


**Figure S1. Biological heterogeneity maps.** The Z-scored maps shown were used to determine regional heterogeneity of local recurrent excitatory $w_i^{EE}$ and excitatory-to-inhibitory $w_i^{EE}$ connectivity weights within the simulations.

T1w/T2w: T1-weighted to T2-weighted ratio, FC G1: principal gradient of functional connectivity, Gene PC1: principal axis of Allen Human Brain Atlas gene expression data, NMDA: N-methyl-D-aspartate receptor density, GABA$_{A/BZ}$: γ-aminobutyric acid type A/Bz receptor density.

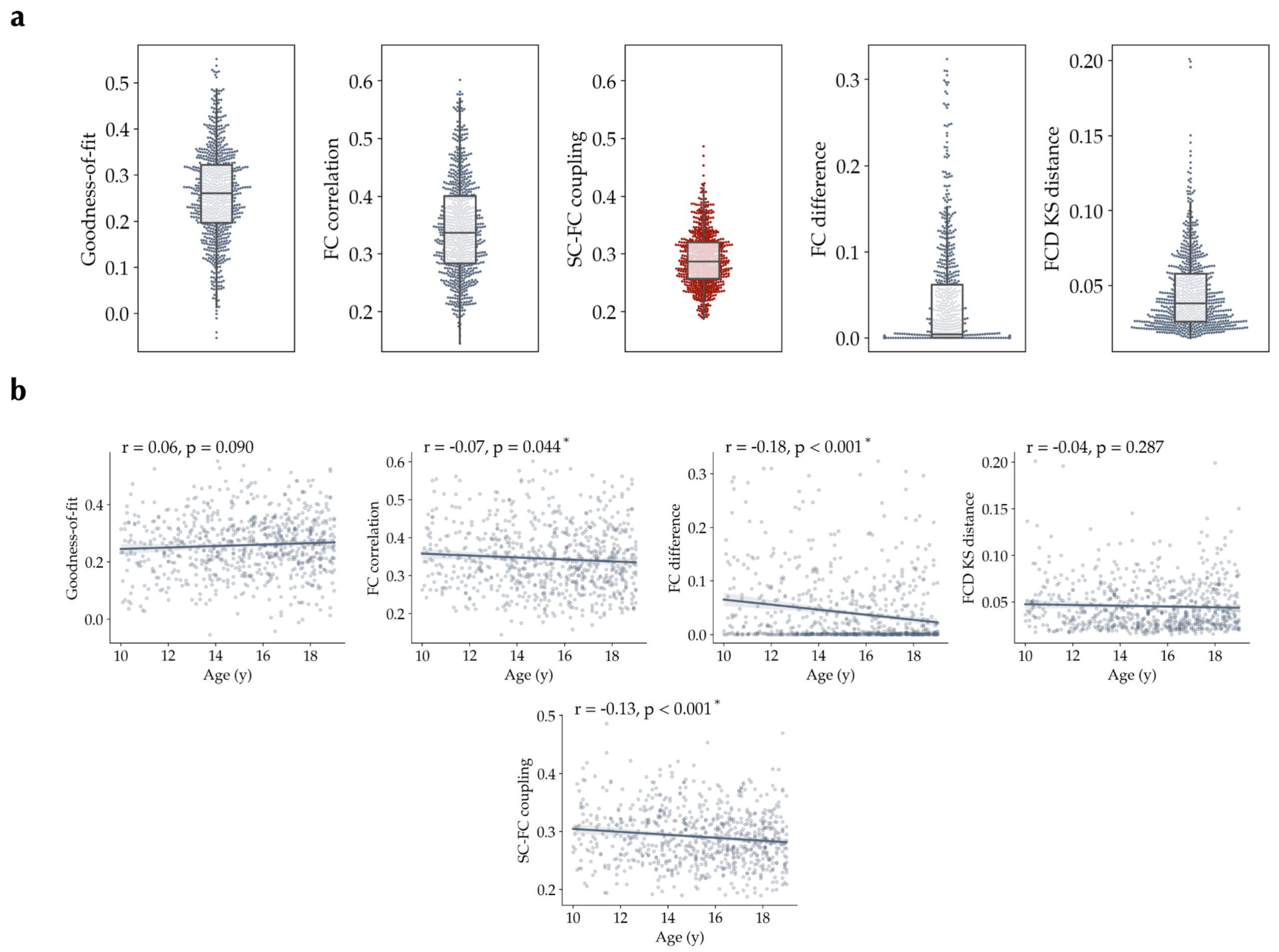


**Figure S2. Goodness-of-fit measures in the PNC dataset. (a)** Distribution of the goodness-of-fit measures. The coupling (correlation) of structural connectome (SC) and empirical functional connectome (FC) is independent of the simulations, and is shown as a reference for FC correlation of simulated and empirical data. **(b)** Pearson correlation of goodness-of-fit measures with age.

FCD: functional connectivity dynamics matrix, KS: Kolmogorov-Smirnov distance.

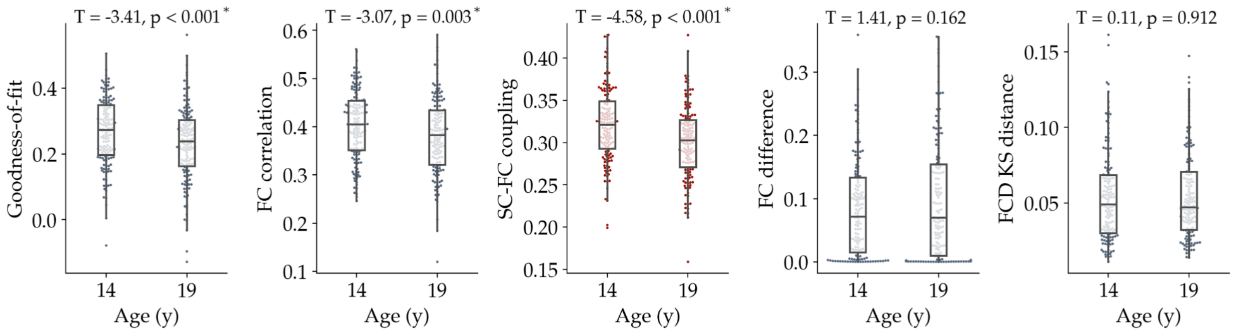


**Figure S3. Goodness-of-fit measures in the IMAGEN dataset.** Distribution of goodness-of-fit measures in the baseline (14 y) and follow-up (19 y) imaging sessions are shown. The goodness-of-fit measures were compared between the two sessions using paired T-test. The coupling of structural connectome (SC) and empirical functional connectome (FC) is independent of the simulations, and is shown as a reference for FC correlation of simulated and empirical data.

FCD: functional connectivity dynamics matrix, KS: Kolmogorov-Smirnov distance.

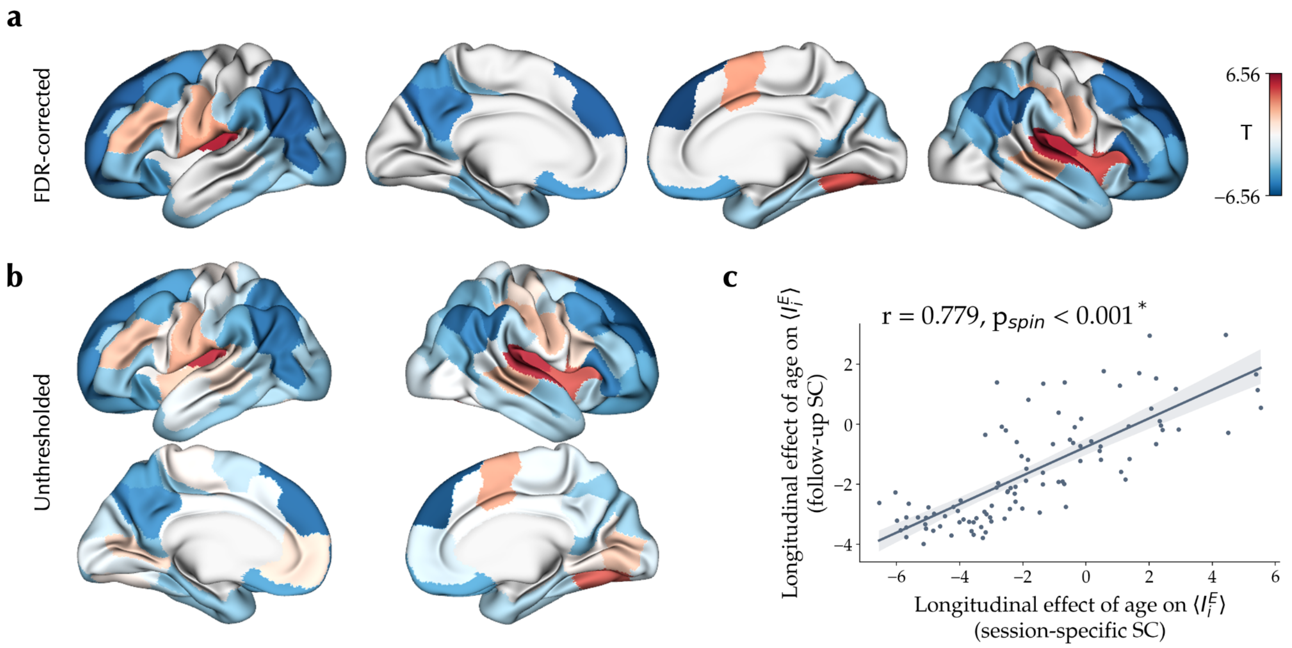


**Figure S4. Longitudinal effect of age on excitation-inhibition balance during adolescence based on session-specific structural connectomes. (a)** Linear effect of age on $\langle I_i^E \rangle$ in a mixed effects model with random intercepts for each subject, after removing outliers and controlling for goodness-of-fit, sex, in-scanner rs-fMRI motion and site, corrected for multiple comparisons using false discovery rate (FDR). **(b)** Unthresholded effect of age on $\langle I_i^E \rangle$. **(c)** Spatial correlation of longitudinal effects of age on $\langle I_i^E \rangle$ using session-specific structural connectomes (SCs) compared to the age effects when the SC of follow-up session was used in the modeling of both sessions.

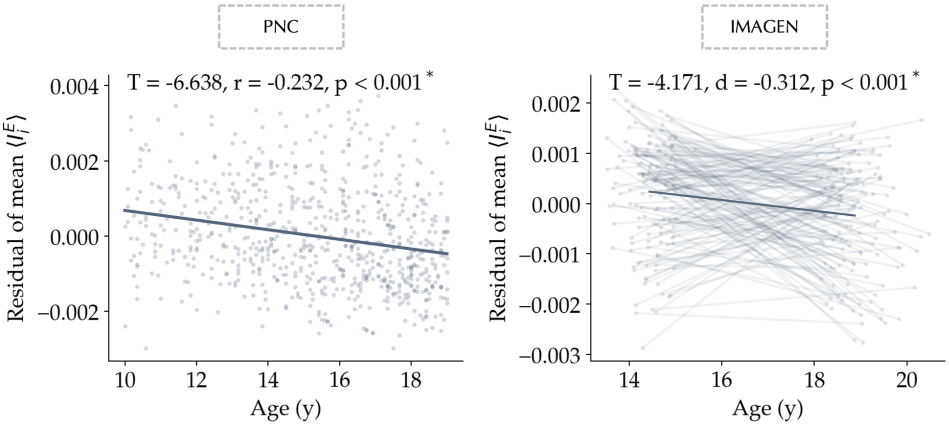


**Figure S5. Effect of age on excitation-inhibition balance across the conjunction mask of significant age effects across datasets.** The effect of age on mean $\langle I_i^E \rangle$ across 33 regions showing significant replicable decreases of $\langle I_i^E \rangle$ in the PNC and IMAGEN (Figure 3c) was investigated controlled for sex, goodness-of-fit, in-scanner rs-fMRI motion, and in IMAGEN, site. Points in both plots represent the residual of mean $\langle I_i^E \rangle$ within the conjunction mask after removing confounds for each individual subject and session. The thick line in PNC shows best linear fit and in IMAGEN connects the average of the two sessions across all subjects. Thin lines in IMAGEN show longitudinal changes of this measure during follow-up in each subject.

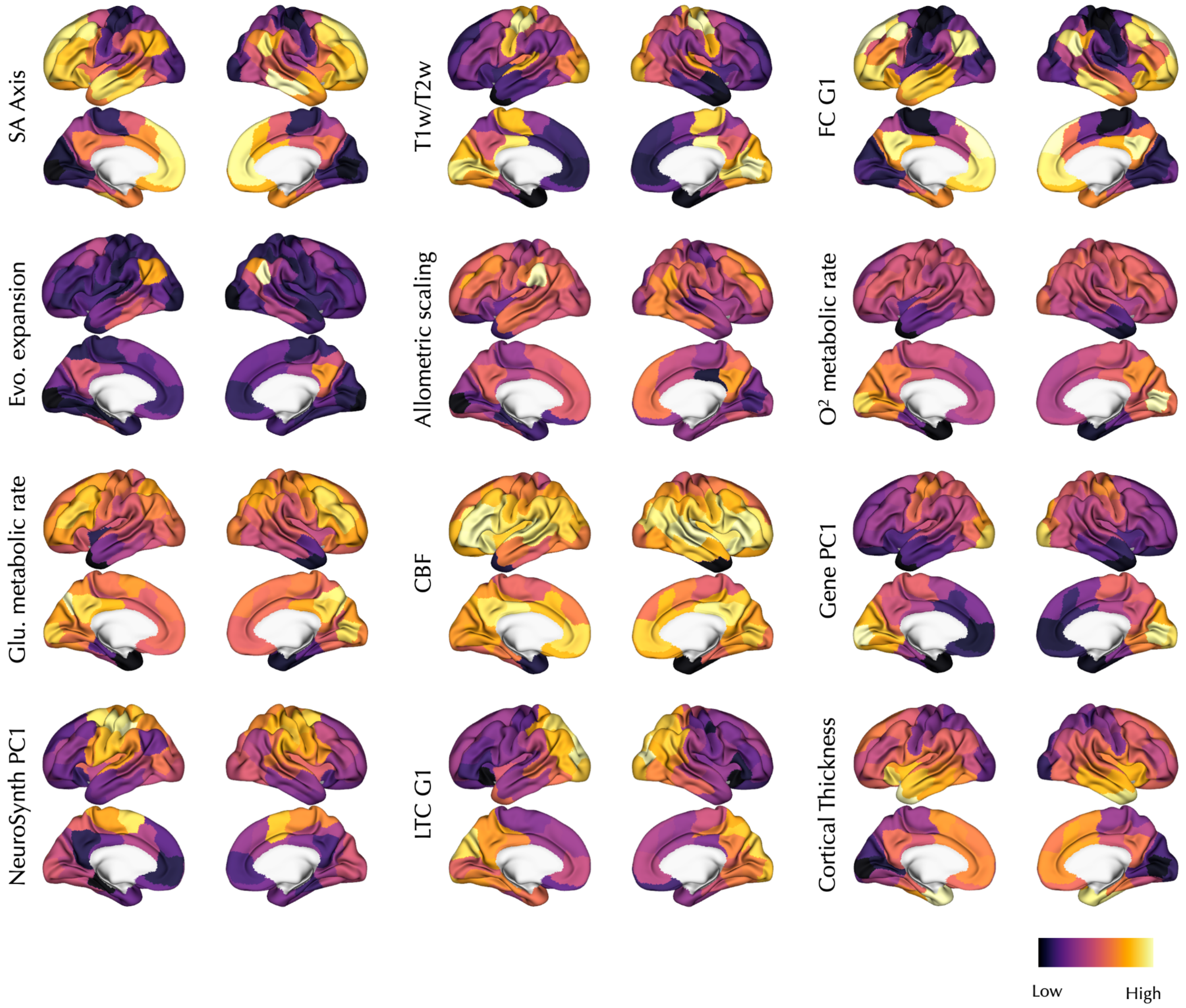


**Figure S6. Multimodal maps of sensorimotor-association cortical axis.** The map of sensorimotor-association (SA) axis proposed in Sydnor et al.[1] and the features it was composed of are shown. See Table S1 for the sources of each map.

T1w/T2w: T1-weighted to T2-weighted ratio, FC G1: principal gradient of functional connectivity, Evo.: evolutionary, CMR: cerebral metabolic rate, Glu.: glucose, CBF: cerebral blood flow, Gene PC1: principal axis of Allen Human Brain Atlas gene expression data, NeuroSynth PC1: Principal component of NeuroSynth meta-analytical maps, LTC G1: principal gradient of laminar thickness covariance.

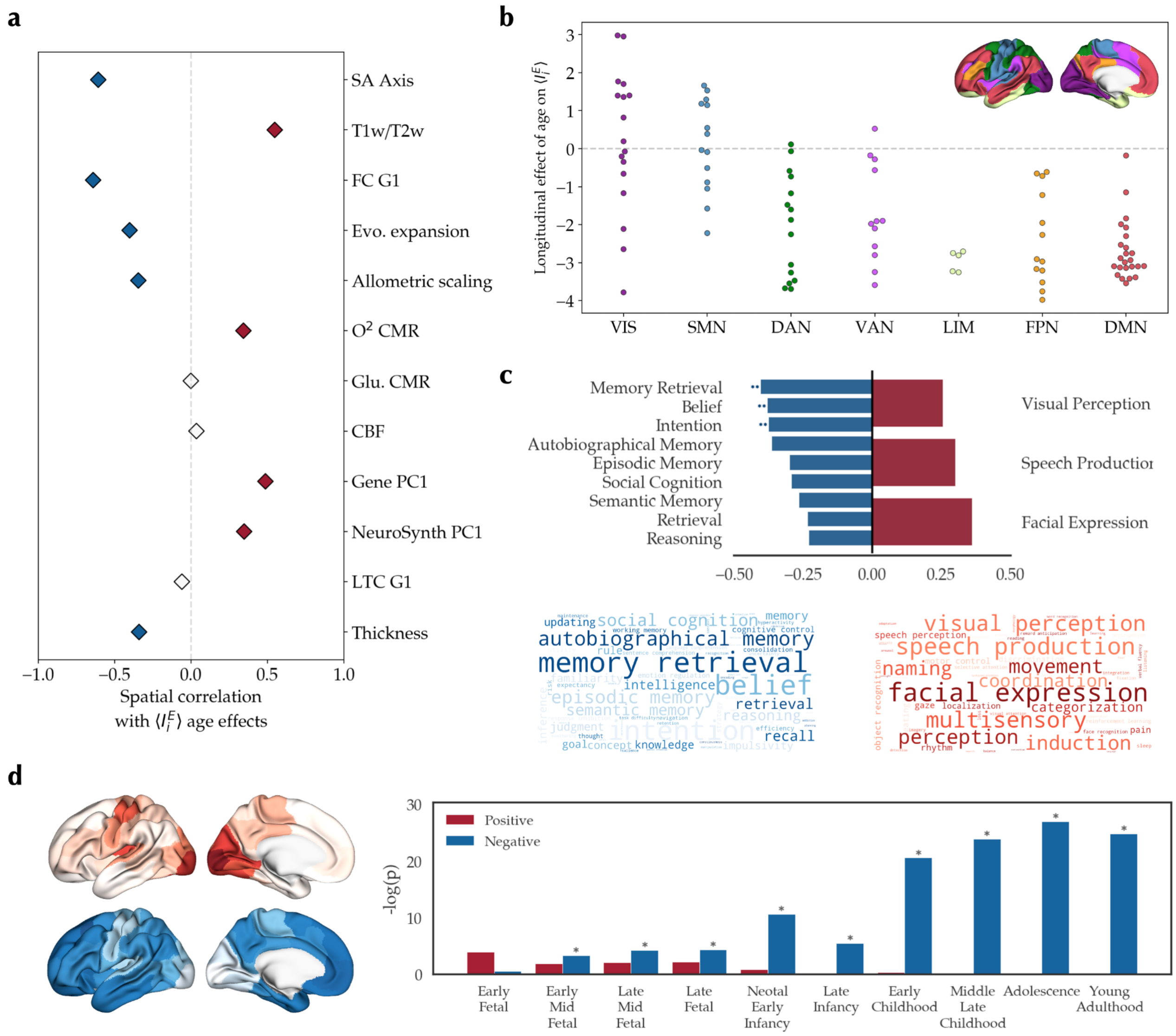


**Figure S7. Embedding of the excitation-inhibition developmental pattern in the IMAGEN dataset along the sensorimotor-association axis.** (**a**) Spatial correlation of $\langle I_i^E \rangle$ age effect in the IMAGEN dataset with maps of sensorimotor-association cortical axis based on Sydnor et al.[1] (Figure S6). Colored diamonds show statistically significant ($p_{spin} < 0.05$) positive (red) and negative (blue) spatial correlations. **(b)** Distribution of $\langle I_i^E \rangle$ age effects across the canonical resting-state networks[2] ($F = 14.57$, $p_{spin} < 0.001$). Post-hoc tests (Bonferroni-corrected) showed significantly more positive age effects in visual (VIS) and somatomotor (SMN) compared to limbic (LIM), dorsal attention (DAN) and default mode networks (DMN) in addition to more positive age effects in SMN compared to ventral attention (VAN) and frontoparietal (FPN) networks. **(c)** *top*: Meta-analytical maps with significant spin correlation to the $\langle I_i^E \rangle$ age effect map. Double asterisks denote associations that were significant after false discovery rate (FDR) adjustment. *bottom*: Word clouds of meta-analytical maps negatively (blue) or positively (red) correlated with the $\langle I_i^E \rangle$ age effect map. Size of the words is weighted by their correlation coefficient. **(d)** *left*: Mean expression of the top 500 genes associated with the $\langle I_i^E \rangle$ age effect map split into sets of negatively- (N = 216, blue) and positively-associated (N = 284, red) genes. *right:* Specific expression analysis of the two sets of genes across developmental stages in the cortex. Y-axis shows the negative log of uncorrected p-values. Asterisks denote significantly enriched developmental stages after FDR adjustment.

SA: sensorimotor-association, T1w/T2w: T1-weighted to T2-weighted ratio, FC G1: principal gradient of functional connectivity, Evo.: evolutionary, CMR: cerebral metabolic rate, Glu.: glucose, CBF: cerebral blood flow, Gene PC1: principal axis of Allen Human Brain Atlas gene expression data, NeuroSynth PC1: Principal component of NeuroSynth meta-analytical maps, LTC G1: principal gradient of laminar thickness covariance.

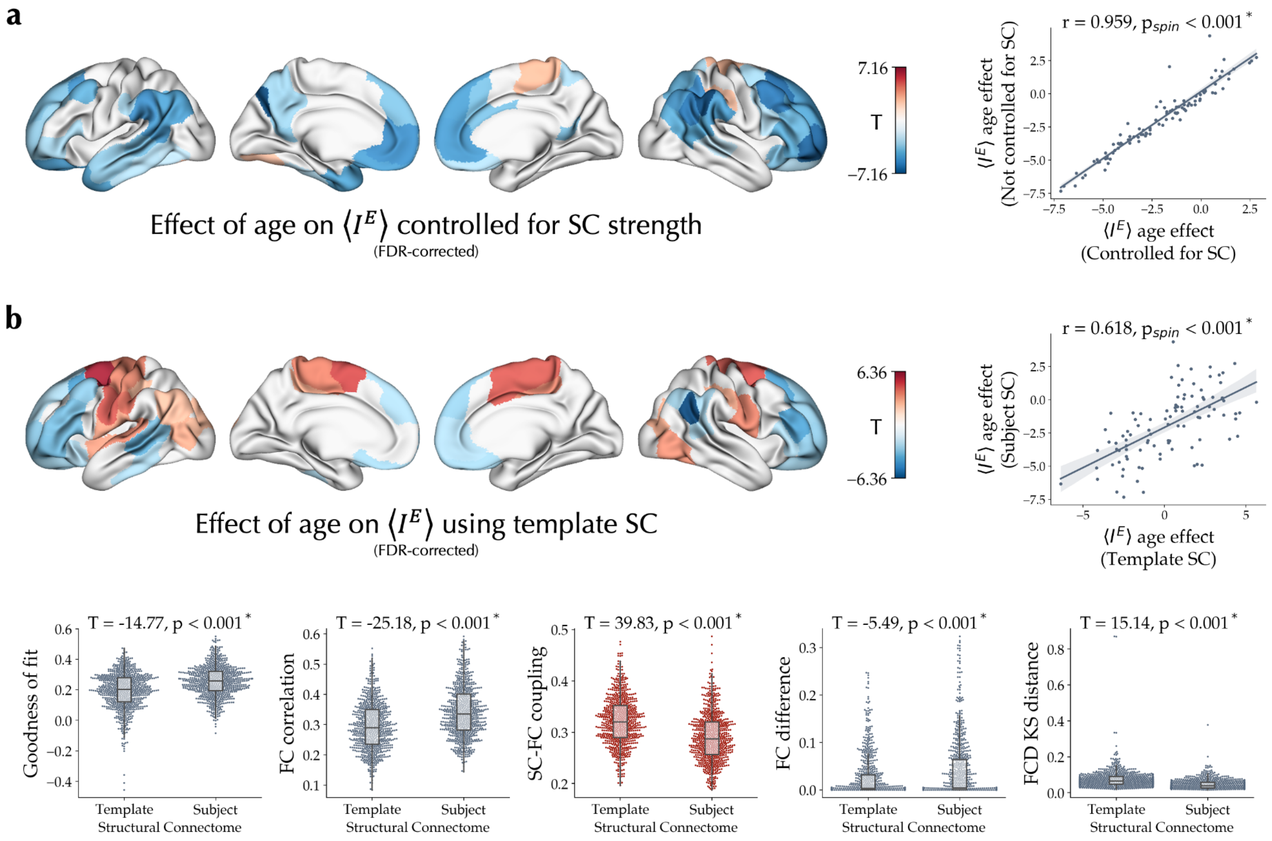


**Figure S8. Robustness of the findings to the effect of inter-individual variability of structural connectome. (a)** *left:* Effect of age on $\langle I_i^E \rangle$ controlled for age, sex, goodness-of-fit, and in-scanner motion, in addition to node-wise structural connectome (SC) strength, that is, row-wise sum of the SC matrix, and corrected for multiple comparisons using false discovery rate (FDR). *right:* Spatial correlation of unthresholded $\langle I_i^E \rangle$ age effect maps with and without controlling for SC strength. **(b)** *top left*: Effect of age on $\langle I_i^E \rangle$ using a fixed template SC in the models, corrected using FDR. *top right:* Spatial correlation of unthresholded $\langle I_i^E \rangle$ age effect maps based on models using subject-specific versus template SCs. *bottom:* Comparison of goodness-of-fit measures and coupling of SC and empirical functional connectome (FC) between models based on subject-specific versus template SCs. T- and p-values resulted from paired T-tests are reported.

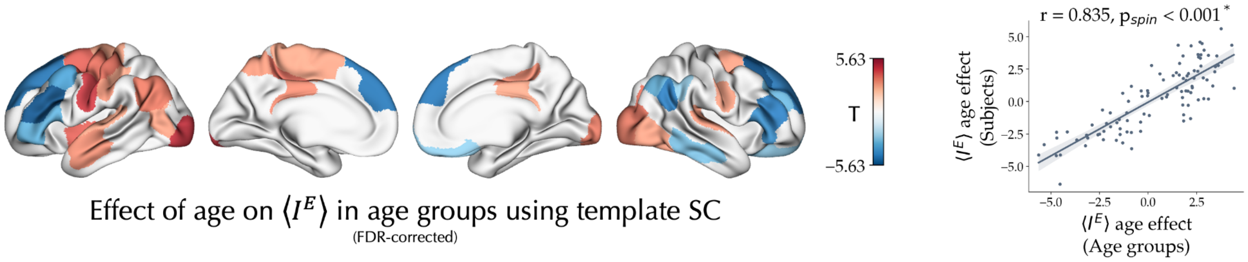


**Figure S9. Effect of age on excitation-inhibition balance in age groups.** *left*: Effect of age on $\langle I_i^E \rangle$ in age groups (N = 30 including 25-27 subject each) using the template SC, after FDR correction. *right:* Spatial correlation of unthresholded $\langle I_i^E \rangle$ age effect maps observed using the template SC at the levels of age groups with the effects observed at the level of subjects (Figure 5a).

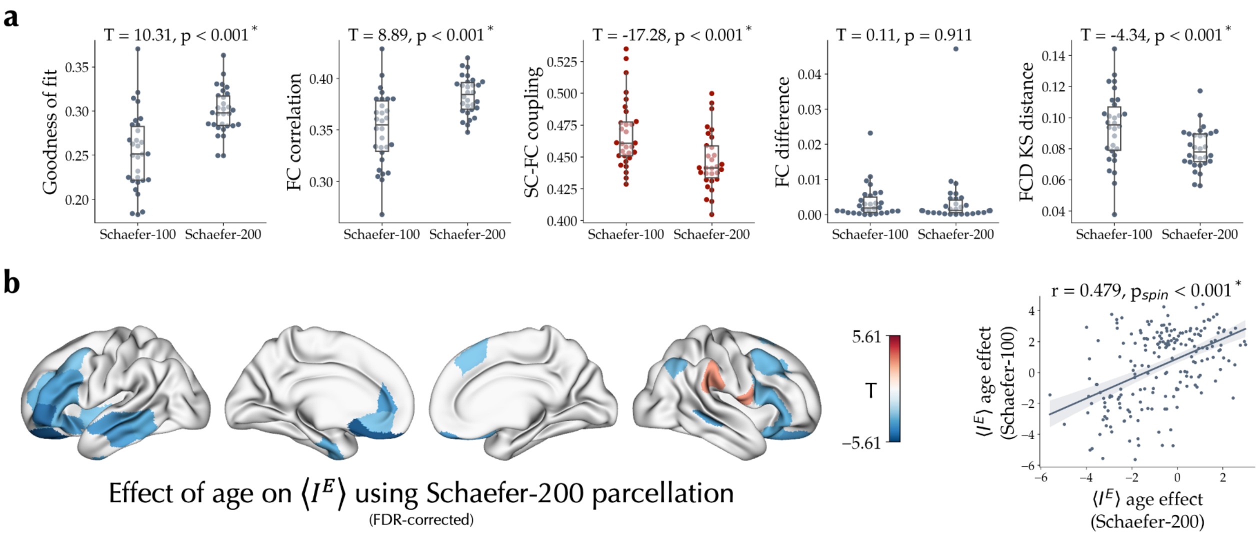


**Figure S10. Robustness of the findings to the effect of parcellation. (a)** Comparison of goodness-of-fit measures and coupling of structural connectome (SC) and empirical functional connectome (FC) between models based on Schaefer-100 and Schaefer-200 parcellations. T- and p-values resulted from paired T-tests are reported. **(b)** *left*: Effect of age on $\langle I_i^E \rangle$ in age groups (N = 30) using Schaefer-200 parcellation, after FDR correction. *right:* Spatial correlation of unthresholded $\langle I_i^E \rangle$ age effect maps between models using Schaefer-100 and Schaefer-200 parcellations.

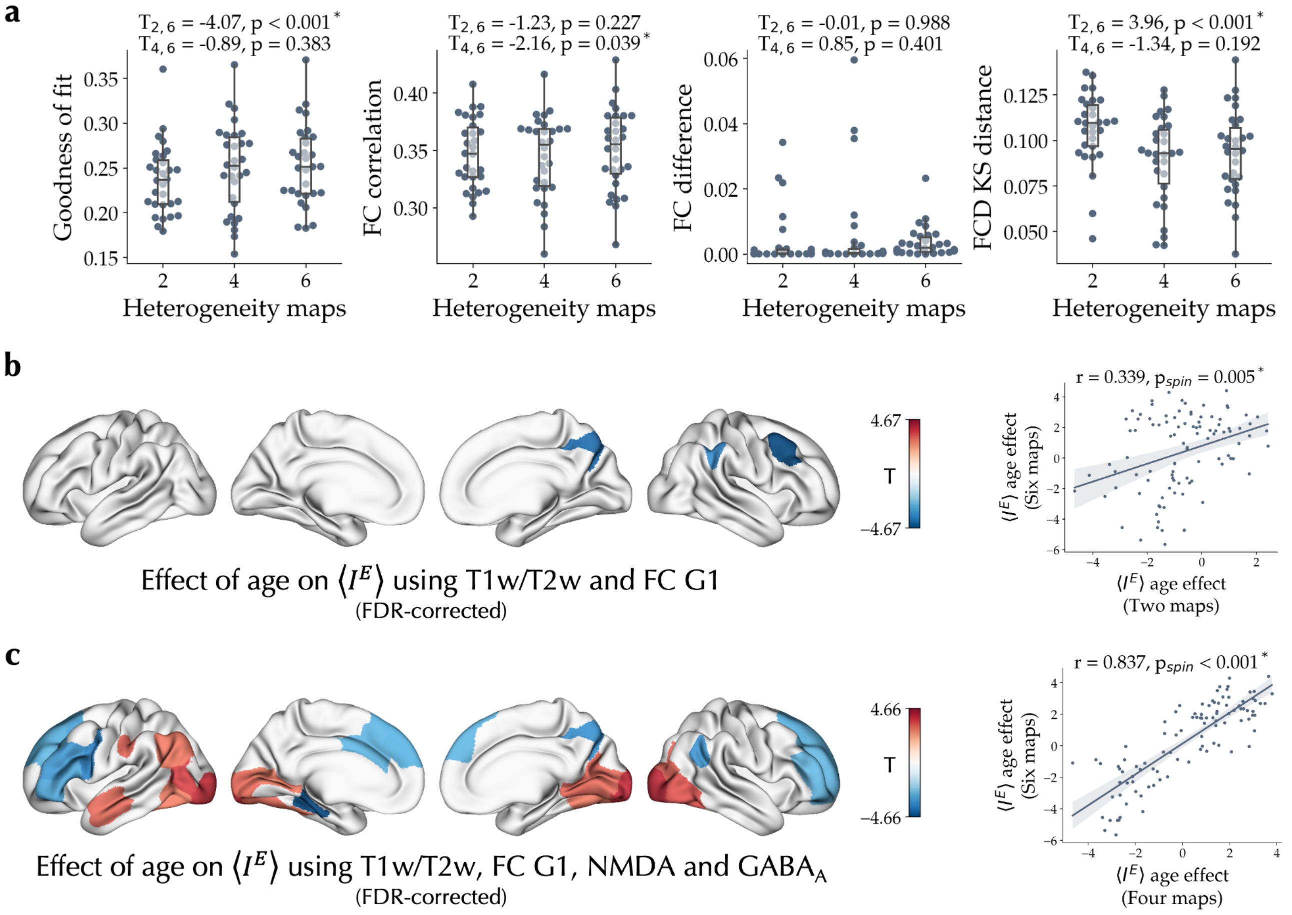


**Figure S11. Robustness of the findings to the effect of heterogeneity maps. (a)** Comparison of goodness-of-fit measures between models based on two, four and six heterogeneity maps. T- and p-values resulted from paired T-tests are reported. **(b)** *left*: Effect of age on $\langle I_i^E \rangle$ in age groups (N = 30) using two heterogeneity maps including T1-weighted to T2-weighted ratio and principal gradient of functional connectivity, after FDR correction. *right:* Spatial correlation of unthresholded $\langle I_i^E \rangle$ age effect maps between models using two and six heterogeneity maps. **(c)** *left*: Effect of age on $\langle I_i^E \rangle$ in age groups (N = 30) using four heterogeneity maps including T1-weighted to T2-weighted ratio, principal gradient of functional connectivity, N-methyl-D-aspartate receptor density and $\gamma$-aminobutyric acid type A/Bz receptor density, after FDR correction. *right:* Spatial correlation of unthresholded $\langle I_i^E \rangle$ age effect maps between models using four and six heterogeneity maps.

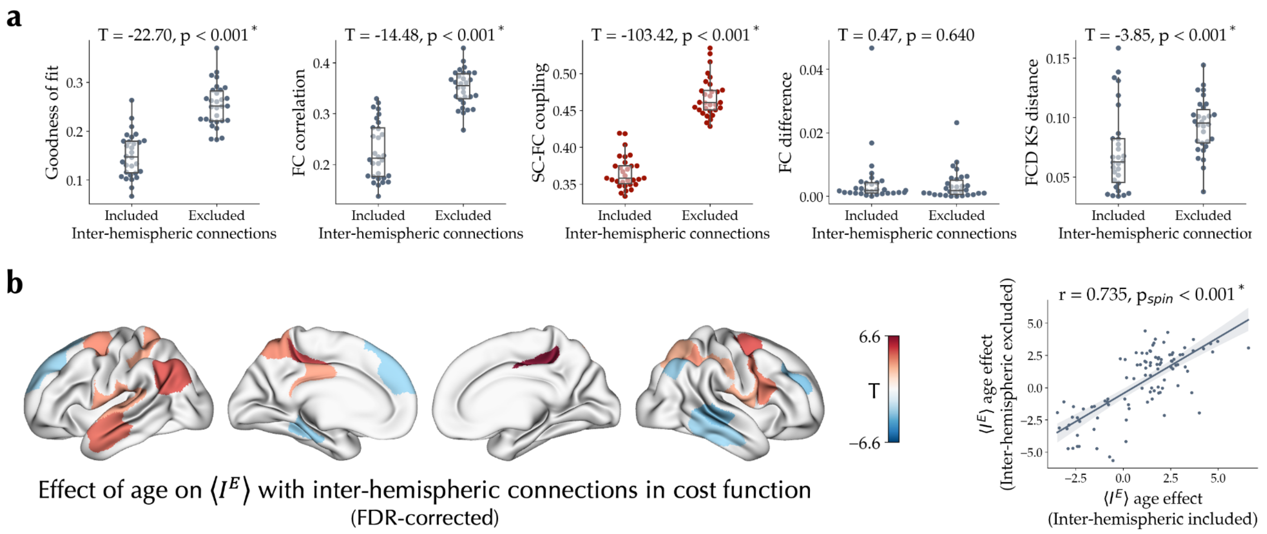


**Figure S12. Robustness of the findings to the inclusion of inter-hemispheric connections in the cost function. (a)** Comparison of goodness-of-fit measures and coupling of structural connectome (SC) and empirical functional connectome (FC) between models fit with the inter-hemispheric connections included or excluded from the cost function. T- and p-values resulted from paired T-tests are reported. **(b)** *left*: Effect of age on $\langle I_i^E \rangle$ in age groups (N = 30) with inter-hemispheric connections included in the cost function, after FDR correction. *right:* Spatial correlation of unthresholded $\langle I_i^E \rangle$ age effect maps between models with the inter-hemispheric connections included or excluded from the cost function.

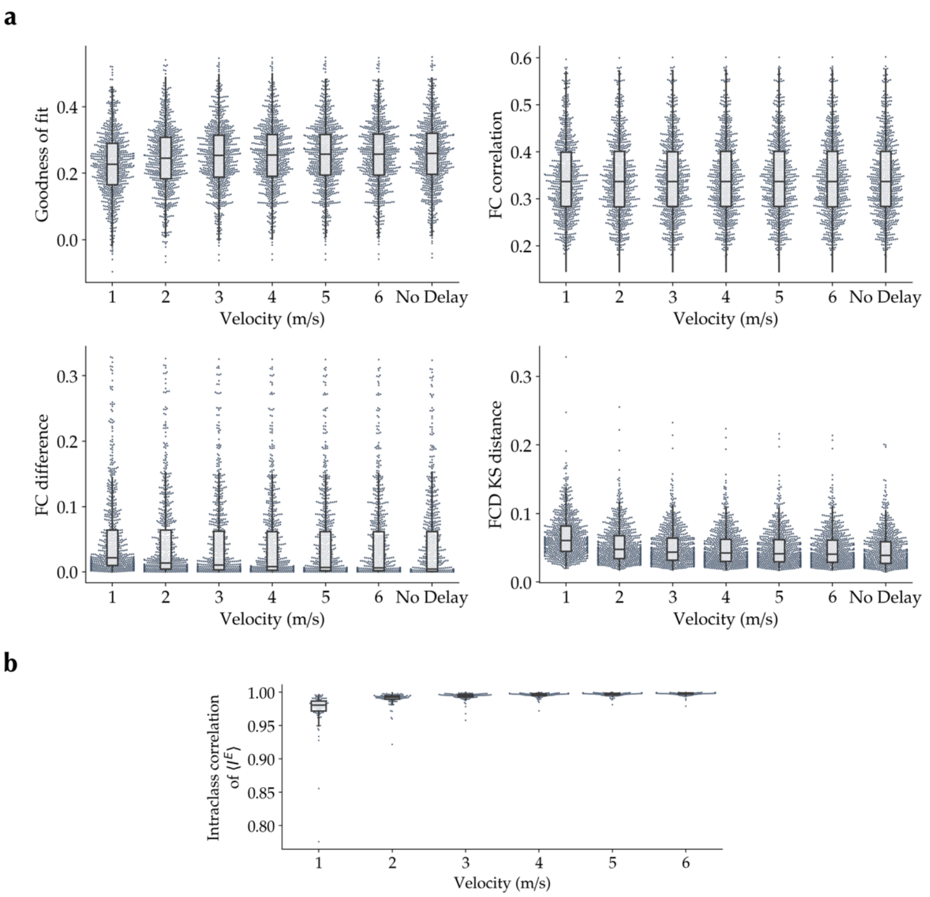


**Figure S13. Effect of conduction delay on goodness-of-fit and $\langle I_i^E \rangle$. (a)** Variation of goodness-of-fit and its components with conduction velocity. The delayed-conduction simulations showed small but significant decreases in GOF, with a mean difference of -0.030±0.020 (T = -41.88, p < 0.001) using a velocity of 1 m/s to a mean difference of -0.003±0.003 (T = -27.76, p < 0.001) using a velocity of 6 m/s. **(b)** Distribution of median absolute deviation intraclass correlation coefficient of $\langle I_i^E \rangle$ between non-delayed-conduction and delayed-conduction simulations across nodes as a function of conduction velocity.

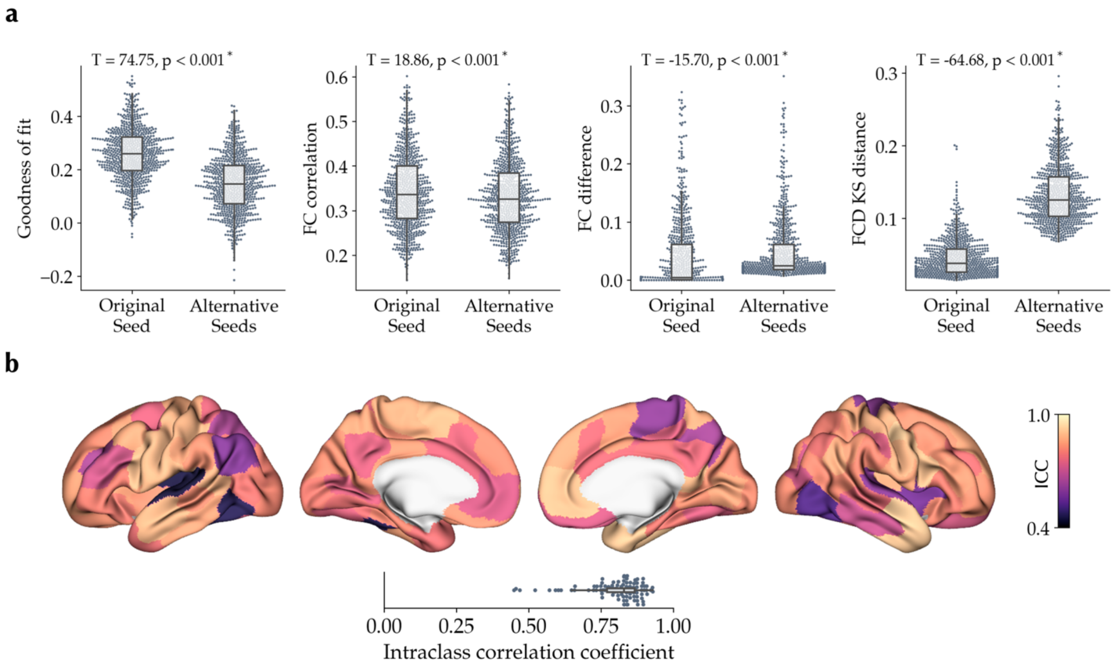


**Figure S14. Effect of Gaussian noise random seed on goodness-of-fit and $\langle I_i^E \rangle$. (a)** Comparison of goodness-of-fit measures between the original simulation and the median of 50 simulations (per subject) using alternative random seeds for the Gaussian noise. T- and p-values resulted from paired T-tests are reported. **(b)** Median of node-wise median absolute deviation intraclass correlation coefficient (ICC) of $\langle I_i^E \rangle$ between the original simulation and the 50 simulations using alternative random seeds for the Gaussian noise.

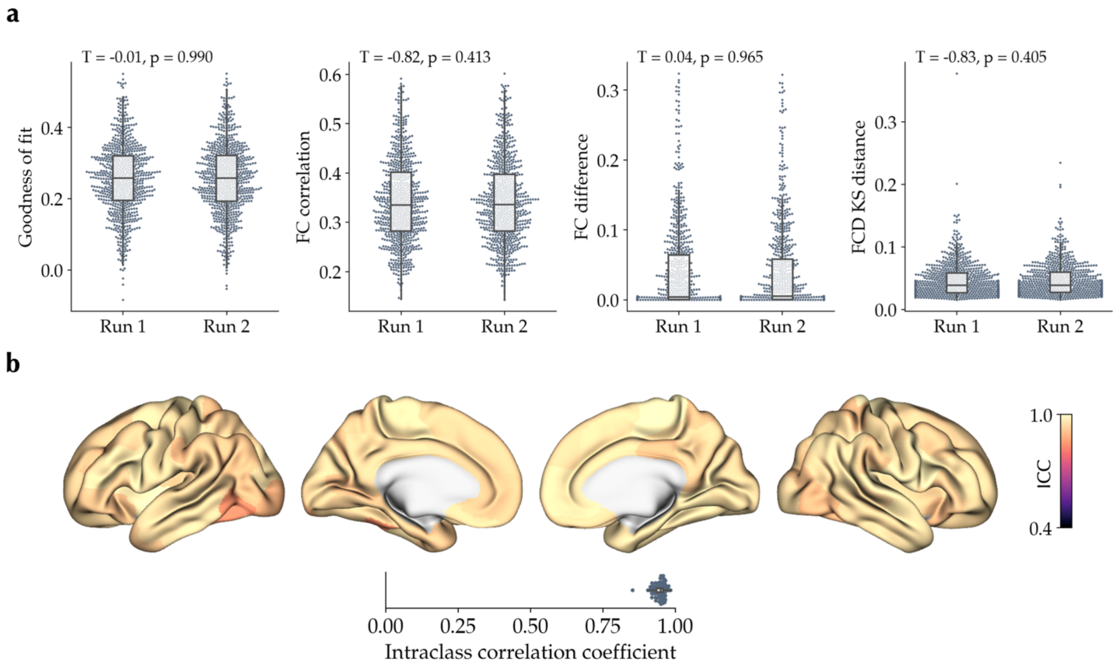


**Figure S15. Effect of optimization seeds on goodness-of-fit and $\langle I_i^E \rangle$. (a)** Comparison of goodness-of-fit measures between the two optimization runs with different random seeds. **(b)** Node-wise median absolute deviation intraclass correlation coefficient (ICC) of $\langle I_i^E \rangle$ between the two runs.

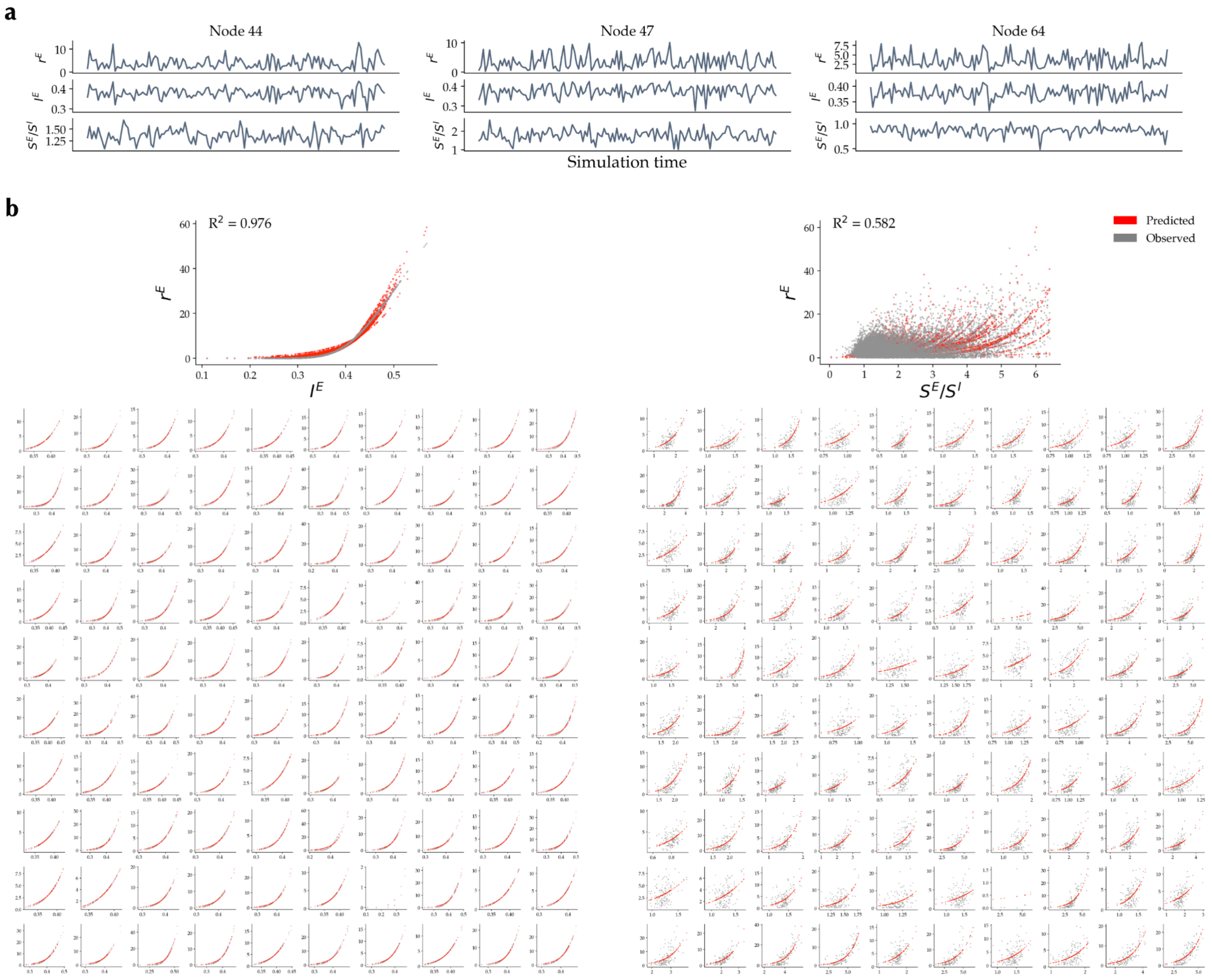


**Figure S16. Association of $r^E$, $I^E$ and $S^E/S^I$ time series in the optimal simulation of an example subject. (a)** Time series of $r^E$, $I^E$ and $S^E/S^I$ in three randomly selected nodes. **(b)** *top*: Association of $r^E$ with $I^E$ (*left*) and $S^E/S^I$ (*right*) across all nodes and time points based on a mixed generalized linear model with an exponential fitting function and random intercepts and slopes per node. Mixed generalized linear model prediction (red) and the observed simulation data (grey) are shown. Note that given $S^E/S^I$ is a ratio and approximates infinity in some data points, we excluded the data points at the top 2.5 percentile of $S^E/S^I$. *bottom*: Model prediction (red) and observed data (grey) for all 100 nodes are shown.

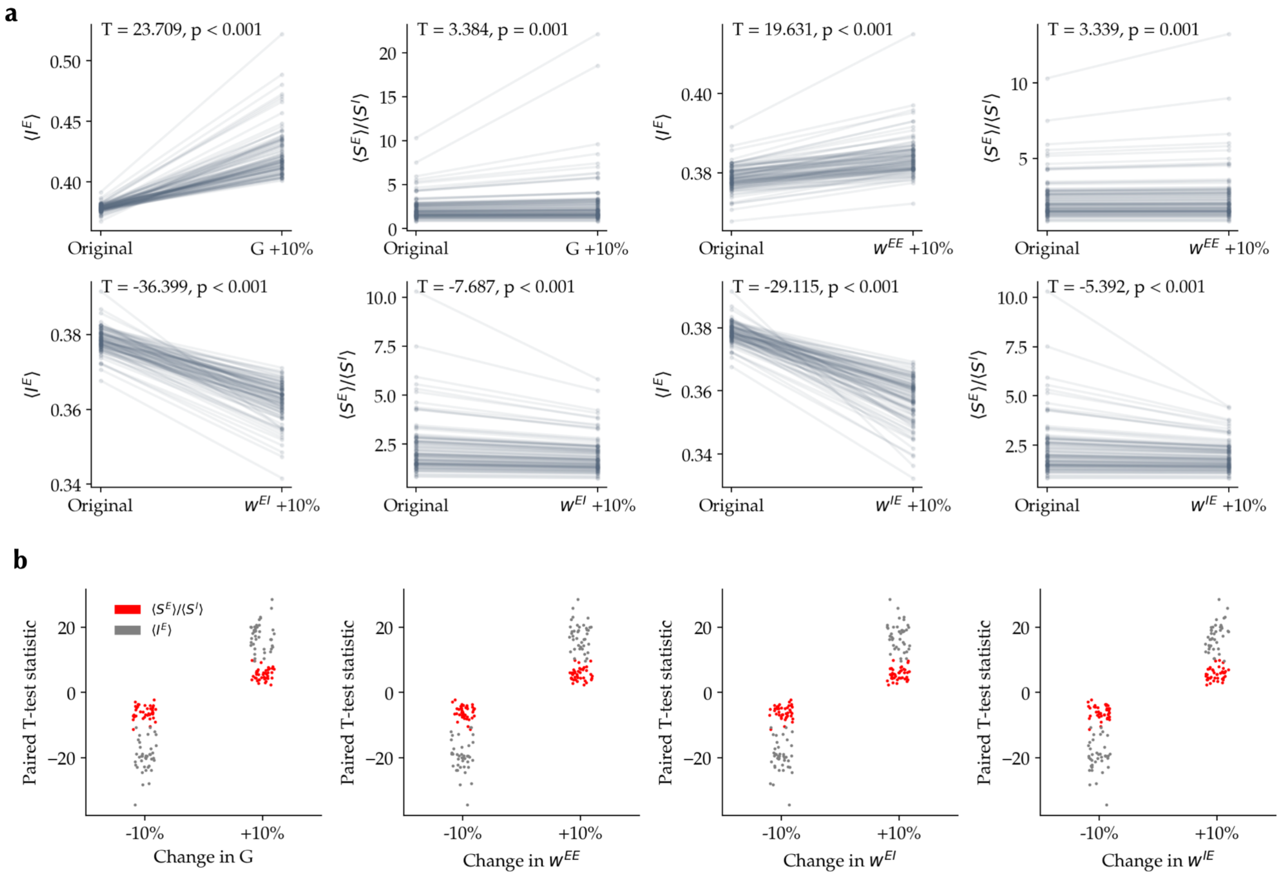


**Figure S17. Effect of model parameters perturbations on $\langle I_i^E \rangle$ and $\langle S_i^E \rangle / \langle S_i^I \rangle$. (a)** Effect of 10% increase in each of the parameters on $\langle I_i^E \rangle$ and $\langle S_i^E \rangle / \langle S_i^I \rangle$ based on the optimal simulation of an example subject ('original' simulation). Paired T-test was used to compare $\langle I_i^E \rangle$ and $\langle S_i^E \rangle / \langle S_i^I \rangle$ across nodes between 'original' and 'perturbed' simulations. **(b)** Paired T statistics comparing $\langle I_i^E \rangle$ (grey) and $\langle S_i^E \rangle / \langle S_i^I \rangle$ (red) between 'original' and 'perturbed' simulations for 40 randomly selected subjects are shown in response to a 10% increase or decrease of each parameter.

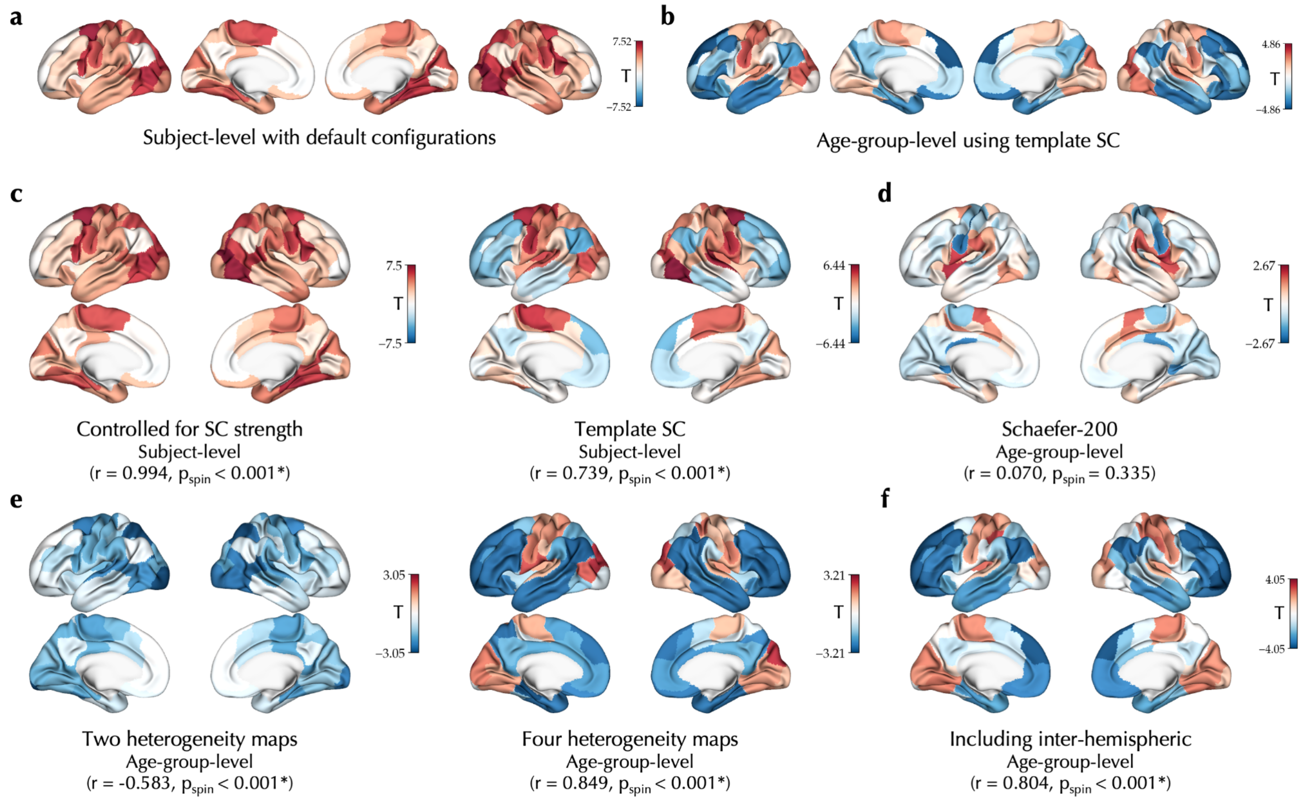


**Figure S18. Effect of age on $\langle S_i^E \rangle / \langle S_i^I \rangle$ and its robustness across different configurations in the PNC dataset.** The unthresholded effect of age on $\langle S_i^E \rangle / \langle S_i^I \rangle$ observed in the PNC dataset using the default configurations at subject level (**a**) or age-group level using template SC (**b**) compared to age effects observed using alternative configurations, including: **(c,** *left***)** additionally controlled for the node-wise structural connectome (SC) strength, that is, row-wise sum of the SC matrix, **(c,** *right***)** using a fixed template SC based on the MICs dataset, **(d)** definition of nodes based on a Schaefer parcellation with higher granularity of 200 nodes, **(e)** using two (*left;* T1-weighted to T2-weighted ratio, principal gradient of functional connectivity) and four (*right;* T1-weighted to T2-weighted ratio, principal gradient of functional connectivity, N-methyl-D-aspartate receptor density, γ-aminobutyric acid type A/Bz receptor density) maps to determine the heterogeneity of regional parameters, and **(f)** by including the inter-hemispheric connections in the goodness-of-fit. In all the panels the statistics indicate spatial correlation of each map with the $\langle S_i^E \rangle / \langle S_i^I \rangle$ age effect observed using the default configurations at subject level (panel **c** correlated with panel **a**) or age-group level (panels **d-f** correlated with panel **b**).

**Table S1. Multimodal maps of sensorimotor-association cortical axis.**

| Map | Reference |
|---|---|
| Sensorimotor-association axis | Sydnor et al.[1] |
| T1-weighted to T2-weighted ratio | Glasser et al.[3,4] |
| Principal gradient of functional connectivity | Margulies et al.[5] |
| Evolutionary expansion | Hill et al.[6] |
| Allometric scaling | Reardon et al.[7] |
| Oxygen cerebral metabolic rate | Vaishnavi et al.[8] |
| Glucose cerebral metabolic rate | Vaishnavi et al.[8] |
| Cerebral blood flow | Satterthwaite et al.[9] |
| Principal component of gene expression data | Hawrylycz et al.[10], Markello et al.[11] |
| Principal component of NeuroSynth | Yarkoni et al.[12], Poldrack et al.[13] |
| Principal gradient of laminar thickness covariance[a] | Saberi et al.[14] |
| Cortical thickness | Glasser et al.[15] |

[a] Substituted instead of externopyramidization map from Paquola et al.[16]. This map was obtained from Saberi et al. while other maps were obtained from neuromaps package[17].

**Table S2. Cognitive and behavioral terms.** Meta-analytical terms of cognitive and behavioral processes based on Hansen et al.[18].

| | | | | |
|---|---|---|---|---|
| action | empathy | listening | response selection | visual perception |
| adaptation | encoding | localization | retention | word recognition |
| addiction | episodic memory | loss | retrieval | working memory |
| anticipation | expectancy | maintenance | reward anticipation | |
| anxiety | expertise | manipulation | rhythm | |
| arousal | extinction | meaning | risk | |
| association | face recognition | memory | rule | |
| attention | facial expression | memory retrieval | salience | |
| autobiographical memory | familiarity | mental imagery | search | |
| balance | fear | monitoring | selective attention | |
| belief | fixation | mood | semantic memory | |
| categorization | focus | morphology | sentence comprehension | |
| cognitive control | gaze | motor control | skill | |
| communication | goal | movement | sleep | |
| competition | hyperactivity | multisensory | social cognition | |
| concept | imagery | naming | spatial attention | |
| consciousness | impulsivity | navigation | speech perception | |
| consolidation | induction | object recognition | speech production | |
| context | inference | pain | strategy | |
| coordination | inhibition | perception | strength | |
| decision | insight | planning | stress | |
| decision making | integration | priming | sustained attention | |
| detection | intelligence | psychosis | task difficulty | |
| discrimination | intention | reading | thought | |
| distraction | interference | reasoning | uncertainty | |
| eating | judgment | recall | updating | |
| efficiency | knowledge | recognition | utility | |
| effort | language | rehearsal | valence | |
| emotion | language comprehension | reinforcement learning | verbal fluency | |
| emotion regulation | learning | response inhibition | visual attention | |
